# Inhibitory Nectin-2/TIGIT interactions between CD163^+^ macrophages and CD8^+^ T cells in spatially defined immune niches associate with poor outcomes in neuroblastoma

**DOI:** 10.64898/2026.10.07.756693

**Authors:** Anne L. Borst, Francisca J. Bergsma, Femke van den Ham, Meggy E.L. Verdonschot, Mojtaba Amini, Arjan Boltjes, Stephanie van Dam, Naima Hiddink Verberne, Noah Bonine, Jeanette H.W. Leusen, Max M. van Noesel, Ronald R. de Krijger, Yvonne Vercoulen, Daniëlle Krijgsman, Judith Wienke

**Author notes:** These authors contributed equally to this work (Anne L. Borst, Francisca J. Bergsma).

## Abstract

**Background:** While immunotherapies have transformed outcomes in several adult cancers, more than half of pediatric patients with high-risk neuroblastoma fail to achieve sustained responses, resulting in poor outcomes. Therefore, understanding the phenotypic and spatial organization of immune cells within the neuroblastoma tumor microenvironment is essential to identify mechanisms driving immunotherapy failure.

**Methods:** To identify spatial determinants of the neuroblastoma immune landscape that may contribute to treatment failure, we dissected immunoregulatory mechanisms at single-cell and protein levels in neuroblastoma samples from 28 patients, using imaging mass cytometry (IMC, n=14) and high-dimensional spectral flow cytometry (n=18).

**Results:** We characterized four distinct cellular neighborhoods, ranging from tumor-dominant to stromal and immune niches. Macrophages were broadly distributed across these niches. In contrast, T and B cells were largely confined to immune niches, forming immature tertiary lymphoid structure-like aggregates within the tumor and stromal cores. T cells displayed a CD69^+^ tissue-resident memory phenotype, with an overall reduced cytotoxic profile (in CD8^+^ T cells) and increased expression of inhibitory receptors (predominantly in CD4^+^ T cells), indicative of dysfunctional features. Intriguingly, T cell functional states differed between spatial niches, with a TCF7^+^/Ki-67^+^ self-renewing profile within immune niches, and increased granzyme B expression in CD8^+^ T cells in tumor cores. Interaction analysis pinpointed CD163^+^ macrophages as the dominant immunosuppressive interaction partner of T cells, particularly within the tumor edge and immune cores. Analysis of inhibitory ligand-receptor interactions identified the TIGIT/PVR/Nectin-2 axis as the main engaged axis between CD163^+^ macrophages and T cells. Nectin-2 expression on macrophages was particularly high within immune cores, and elevated in macrophages with direct T cell contact, suggesting a high probability of *in situ* engagement. Consistent with this immunosuppressive role, both CD163 and Nectin-2 expression on macrophages correlated with poor patient survival, substantiating the clinical relevance of inhibitory macrophage-T cell interactions.

**Conclusions:** This study reveals T cell-excluded tumor niches and targetable inhibitory interactions between CD163^+^ macrophages and T cells. These insights provide a rationale for combinatorial immunotherapeutic strategies that both enhance T cell infiltration into tumor-rich areas and disrupt inhibitory signaling to improve immunotherapy efficacy in neuroblastoma.

## INTRODUCTION

Immunotherapies have revolutionized the treatment of adult cancers, but their efficacy in pediatric solid tumors remains limited^1,2^. A deeper understanding of the immunoregulatory mechanisms within the tumor microenvironment (TME) of pediatric tumors is essential to identify new immunotherapeutic targets and enhance the efficacy of immunotherapies.

Around 10% of all pediatric cancer deaths are caused by neuroblastoma, the most common extracranial solid tumor in children^3^. It originates from the sympathetic nervous system and can present in various risk groups with distinct prognoses and treatment strategies, based on molecular factors and disease presentation, including patient age, tumor histology, localized versus metastatic disease, and genomic alterations. Among this, *MYCN* amplification is an important driver of poor prognosis^4,5^. While patients with localized low-risk tumors have a good prognosis with >90% survival, the 5-year survival rate of patients with widespread high-risk disease is still below 60% due to relapses^6^, despite receiving an intensive treatment protocol that includes induction chemotherapy, surgery, high-dose consolidation chemotherapy, autologous stem cell transplantation, radiotherapy, and anti-GD2 immunotherapy^7^. Although the introduction of anti-GD2 into the standard of care improved survival rates of high-risk patients by approximately 15%^6,8^, the prognosis of high-risk patients is still dismal. This underscores the urgent need for improved therapies for high-risk neuroblastoma patients.

The promising results of immunotherapy with anti-GD2 antibodies in neuroblastoma, as well as the strong association between the extent of lymphocyte infiltration and patient prognosis^9–12^, have sparked the development of additional, T cell-based immunotherapies, including GD2 and B7-H3 targeting chimeric antigen receptor (CAR) T-cell therapy and immune checkpoint inhibition^13–16^. However, survival benefits from most immunotherapies are still modest, since clinical responses do not persist in the majority of patients.

This is likely due to several challenges that neuroblastomas present. First, neuroblastomas are poorly recognized by the immune system due to a low mutational burden and low expression of MHC-I^17,18^. *MYCN*-amplified tumors additionally display reduced immune cell infiltration compared to non-amplified tumors^12^. Second, even when T cells do infiltrate the tumor, their function may be hampered by immunoregulatory networks within the TME, as shown in recent studies utilizing single-cell RNA sequencing (scRNAseq)^19–22^. For instance, T cell reactivity has been reported to be hampered by immunosuppressive cells, including pro-tumorigenic macrophages and regulatory T cells (Tregs), by expression of immune checkpoints such as PD-1 and TIGIT^22,23^, and by inhibitory soluble mediators including MIF^24^.

Despite the valuable insights that these studies provided, it has been established that scRNAseq studies typically suffer from data sparsity^25^, and that RNA and protein levels do not always correlate^26^. This warrants a detailed protein-based investigation into the immune interactions in neuroblastoma. Most importantly, current data lack the spatial context of the neuroblastoma immune landscape, while recent studies have demonstrated that the spatial context of immune infiltration and immune interactions is a key determinant of immunotherapy efficacy^27,28^.

Therefore, we employed high-dimensional spectral flow cytometry and imaging mass cytometry (IMC) to gain detailed insights into the (spatial) immune landscape of neuroblastoma, including the presence of immunosuppressive cells and immunosuppressive axes at the single-cell and protein levels, to guide the development of more effective immunotherapies for high-risk neuroblastoma. We observed a spatially structured TME in which dysfunctional T cells are excluded from tumor cores and sequestered within immune cores. Moreover, we identified inhibitory interactions, including the Nectin-2/TIGIT axis, with neighboring CD163^+^ immunosuppressive macrophages, which was prognostic for patient survival.

## MATERIAL AND METHODS

### Patients and tissue processing

From the Princess Máxima Center (Utrecht, the Netherlands), 18 patients with neuroblastoma were included between January 2020 and January 2022. Samples were obtained by surgical resection after induction chemotherapy (n=35). Clinical data was obtained retrospectively on 11-06-2026 (**Suppl. Table 2**). Written informed consent for this study was obtained for all patients, from themselves, parents or guardians. Healthy adult donor blood was obtained from the shared healthy donor blood biobank from the University Medical Center Utrecht (biobank number 18-774; written informed consent was obtained from all donors). This study received approval by the ethical committee as per national legislation. Tumor samples were obtained through an institutionally approved research study by the institutional review board of the Erasmus Medical Center Rotterdam (MEC-2016-739) and the biobank committee of the Princess Máxima Center (PMCLAB2020.093). This study was conducted in accordance with the guidelines of the Declaration of Helsinki.

For spectral flow cytometry, fresh tumor tissue was minced into pieces of ∼1mm^3^ and dissolved using collagenase IV for max 1 hour at 37°C, filtered through a 70µm cell strainer and frozen in 10% DMSO until further use. Peripheral blood mononuclear cells (PBMCs) were freshly isolated from healthy adult blood by Ficoll-Paque density gradient centrifugation. Cells were frozen in 10% DMSO until further use.

For IMC, five consecutive formalin-fixed paraffin-embedded (FFPE) tissue sections of 4µm were prepared: the middle section was stained with hematoxylin-eosin (HE), with the adjacent sections used for the myeloid-and lymphoid-focused IMC panels. The outermost sections were stained by immunohistochemistry (IHC) for CD15 (granulocytes) and CD3 (T cells), respectively, using DAB detection to guide region of interest (ROI) selection by an expert pathologist (RRdK), who selected two ROIs of 1000×1000 µm^2^ per sample targeting tumor regions with immune infiltrate.

### Spectral flow cytometry staining

Cryopreserved tumor digests were thawed, washed twice with PBS, and stained with Fixable Viability Dye eFluor™ 506 (ThermoFisher, 65-0866-14) for 20 min at 4°C. Cells were subsequently washed with PBS, stained with fluorescently labelled surface antibodies (antibody panels in Suppl. Table 1) for 20 min at 4°C, washed with staining buffer (PBS with 2% FCS and 2mM EDTA) and fixed and permeabilized using the FOXP3/Transcription factor staining buffer set (Invitrogen, 00-5523-00). Lastly, cells were stained with intracellular antibodies (**Suppl. Table 1**) for 30 min at 4°C, washed and acquired on an Aurora flow cytometer (Cytek Biosciences).

### Spectral flow cytometry analysis

Data were analyzed using both manual and computational methods. Data were exported from FlowJo and used for unsupervised clustering using the R package cyCONDOR (version 0.2.1)^29^. Characteristics of each cluster were evaluated using marker expression in UMAP space and in heatmaps. Cells were gated manually in FlowJo version 10.10 to identify major cell populations. Marker expression by each cell population was extracted as percentage marker expressing cells and mean fluorescent intensity (MFI). For analysis of marker expression, populations with <20 cells were excluded. Boxplots were generated with R package ggplot2 (version 3.5.1) and barplots with GraphPad Prism (GraphPad Software). Heatmaps were used to visualize marker expression patterns and were generated using the heatmap() function in base R. To visualize marker expression patterns in a heatmap, marker MFI’s were hierarchically clustered based on Euclidean distance and Ward’s method. For correlation analyses, spearman correlation coefficients were calculated. To identify possible immune checkpoint interactions, we calculated the interaction probability with the following formula: log(1+(R_MFI_ * R_%_ x L_MFI_ * L_%_ x n_A_ * n_B_ / n^2^ x (1/K_D_))), in which ‘R’ stands for receptor, ‘L’ for ligand, ‘n’ for number of cells of cell type A and B and K_D_ for dissociation constant. We normalized the MFI of the receptor (R_MFI_) expressed by cell type A and ligand (L_MFI_) expressed by cell type B to range from 0 to 1, which was calculated by: (MFI – MFI_min_) / (MFI_max_ – MFI_min_). The expression of the receptor by cell type A or ligand by cell type B was calculated by multiplying the normalized MFI with the percentage of receptor or ligand expressing cells of cell type A or B, respectively (R_%_ or L_%_). To account for the population size, the number of cells of type A (n_A_) and B (n_B_) were multiplied and divided by the squared total number of cells. The binding affinity was defined by the dissociation constant (1/K_D_), which was derived from literature^30–34^.

### IMC antibody optimization and tissue staining

Carrier-free antibodies for the myeloid-and lymphoid-focused panels were conjugated with metal isotopes using the Maxpar X8 Labeling Kit (Standard BioTools) and stored in phosphate-buffered saline until use (**Supplementary Table 1**). Antibodies were validated on control tissues (colon, inflamed colon, tonsil, and placenta) by immunofluorescence (IF) for specificity and titrated by IMC on these control tissues to determine optimal dilutions.

FFPE sections were deparaffinized, antigen retrieved (target retrieval solution pH 9.0) and blocked with SuperBlock buffer. Tissue sections were then stained as previously described^35,36^. Briefly, sections were incubated overnight at 4°C with anti-SIRPα (myeloid panel) or anti-CTLA-4 (panel 1) in Tris-buffered saline with 0.1% Tween® (TBST) containing 0.5% Bovine Serum Albumin (BSA). After washing, sections were stained with secondary antibodies rabbit-161Dy (SIRPα) or rabbit-154Sm (CTLA-4) in TBST containing 0.5% BSA for 1 hour at RT. After washing, sections were exposed to the metal-labelled antibody cocktails in TBST containing 0.5% BSA for overnight incubation at 4°C. After washing, the sections were stained with DNA intercalator Ir193 (Standard BioTools, 1:500) and DAPI (Sigma-Aldrich, 1:10.000) for 1h at RT. Following fluorescence microscopy, slides were processed for IMC acquisition on the Hyperion Imaging System (Standard BioTools), as previously described^36^.

### IMC data processing

The raw IMC images (.mcd files) were converted to TIFF format using the MCD viewer (Standard BioTools) and image channels were normalized using Percentile Normalization GUI Image denoising (PENGUIN)^37^. Optimal percentiles and thresholds were chosen for each channel separately by manual inspection of several images and applied to every ROI. Denoised images were segmented in CellProfiler (v4.2.8) by first generating nuclear masks using the Cellpose-SAM algorithm^38^ on the H3 histone channel, followed by expansion of these nuclear masks using membrane markers for lymphoid cells (CD20, CD3, CD8), myeloid cells (CD11c, CD68), stromal cells (aSMA) and tumor cells (CD56, Synaptophysin), with additional CD14 for the myeloid segmentation in panel 2 and CHGA for the tumor cell segmentation in panel 1. Next, the generated masks and denoised images were loaded into ImaCytE^39^ to extract the single-cell data as.fcs file per sample, which was subsequently merged with the cell metrics generated from CellProfiler and the patient metadata to generate a comprehensive data frame containing spatial coordinates and marker intensities for downstream analysis in R (v4.4.0).

To enable robust downstream analysis, cells with gadolinium residue, indicative of MRI contrast agent contamination, as described previously^40^, were identified based on simultaneous high expression (above the 95^th^ percentile) of the four gadolinium-conjugated markers in each panel (panel 1: CD127, FOXP3, TIGIT, TCRγδ; panel 2: FOXP3, CD4, IL10, pSTAT3) and excluded from downstream analysis. Additionally, cells with a coefficient variation (CV) of zero across all lineage markers were excluded as low-quality cells. Lineage markers used for CV calculation included CD3, CD8, CD4, CD45, CD20, CD68, CD11c, aSMA, CD31, Synaptophysin, CHGA and CD56 for panel 1 and CD3, CD8, CD4, CD45, CD20, CD68, CD11c, CD11b, CD15, CD14, aSMA, CD31, Synaptophysin, and CD56 for panel 2. Furthermore, one patient (M313AAB) was excluded from all analyses as the tumor was retrospectively reclassified by the pathologist as healthy adrenal gland. One patient (M223AAD) was excluded from analysis due to insufficient cell segmentation resulting from densely packed nuclei within a tertiary lymphoid-like structure (TLS). Lastly, one patient (M050AAB) was excluded from panel 1 analyses due to poor staining quality. These quality steps yielded a total of 177.813 (panel 2) and 166.555 (panel 1) cells for downstream data analysis.

### IMC data analysis

Cell phenotypes were assigned by supervised classification based on the expression of specific lineage makers, as described in **Supplementary Figure 1A**. Cellular neighborhood (CN) and cell-cell interaction analyses were performed using the workflow proposed by Windhager et al.^41^ using the R package *imcRtools* (v1.17.0). For each ROI, an expansion graph was constructed with an expansion distance of 30 µm around each cell. Neighboring cell type compositions were subsequently aggregated per cell using the *aggregateNeighbors* function based on the constructed expansion interaction graph and cells were clustered into 4 CNs. Cell-cell interactions across all ROIs and CNs were assessed using the *testInteractions* function applying the conditional permutation-based method with p threshold of 0.05.

For the spatial CD8 T cell states, adapted from Meyer et al.^42^, each CD8 T cell was assigned to one of the three states based on the proportion of tumor cell neighbors in the aggregated expansion interaction graph: isolated (0% tumor cell neighbors), bystander (less than 50% tumor cell neighbors), and infiltrating (50% or more tumor cell neighbors). Each patient was subsequently assigned to a spatial tumor phenotype group based on the predominant CD8 T cell state: cold, excluded, or inflamed. For all marker expression analyses, ROIs of patients containing fewer than 10 cells of the relevant cell type were excluded from analysis.

### Statistical analysis

Statistics for flow cytometry were performed using GraphPad Prism and R. Comparisons between two groups were made by Wilcoxon rank-sum test with Benjamini-Hochberg (BH) correction applied for multiple testing. Markers with padj<0.05 and a fold change of >0.6 or <-0.6 were considered differentially expressed and visualized with volcano plots, generated with ggplot. Comparisons between 3 or more groups were made by ANOVA with Sidak or Tukey post hoc test for multiple comparisons, as indicated in the figure legends.

Statistical analyses for IMC were performed using the Wilcoxon rank-sum test for comparisons between two groups. Comparison across multiple groups were assessed using pairwise Wilcoxon rank-sum test with Benjamini-Hochberg (BH) correction applied for multiple testing. Differences in cellular composition were assessed using the propeller (limma-based differential proportion analysis) method from the *speckle* (v1.4.0) package. For Kaplan-Meier survival analyses, groups were dichotomized by median and relapse samples were excluded. Adjusted p-values <0.05 were considered statistically significant.

## RESULTS

### Lymphoid and myeloid cells display distinct spatial patterning in the neuroblastoma TME

To comprehensively characterize the TME of neuroblastoma spatially and at the protein level, we performed spectral flow cytometry and IMC on tumor samples obtained by surgical resection after induction chemotherapy. For each technique, dedicated lymphoid (Panel 1) and myeloid/tumor (Panel 2) antibody panels were designed to enable in-depth phenotypic and spatial characterization of infiltrating immune cells and tumor cells (**Table S1**). Spectral flow cytometry was performed on samples from 18 patients and IMC on samples from 14 patients (**Figure 1A and Table S2**). 22 patients had high-risk neuroblastoma, five patients medium-risk and one patient low-risk. The clinical characteristics of the 28 included patients are summarized in **Supplementary Table S2**.

**Figure 1.**
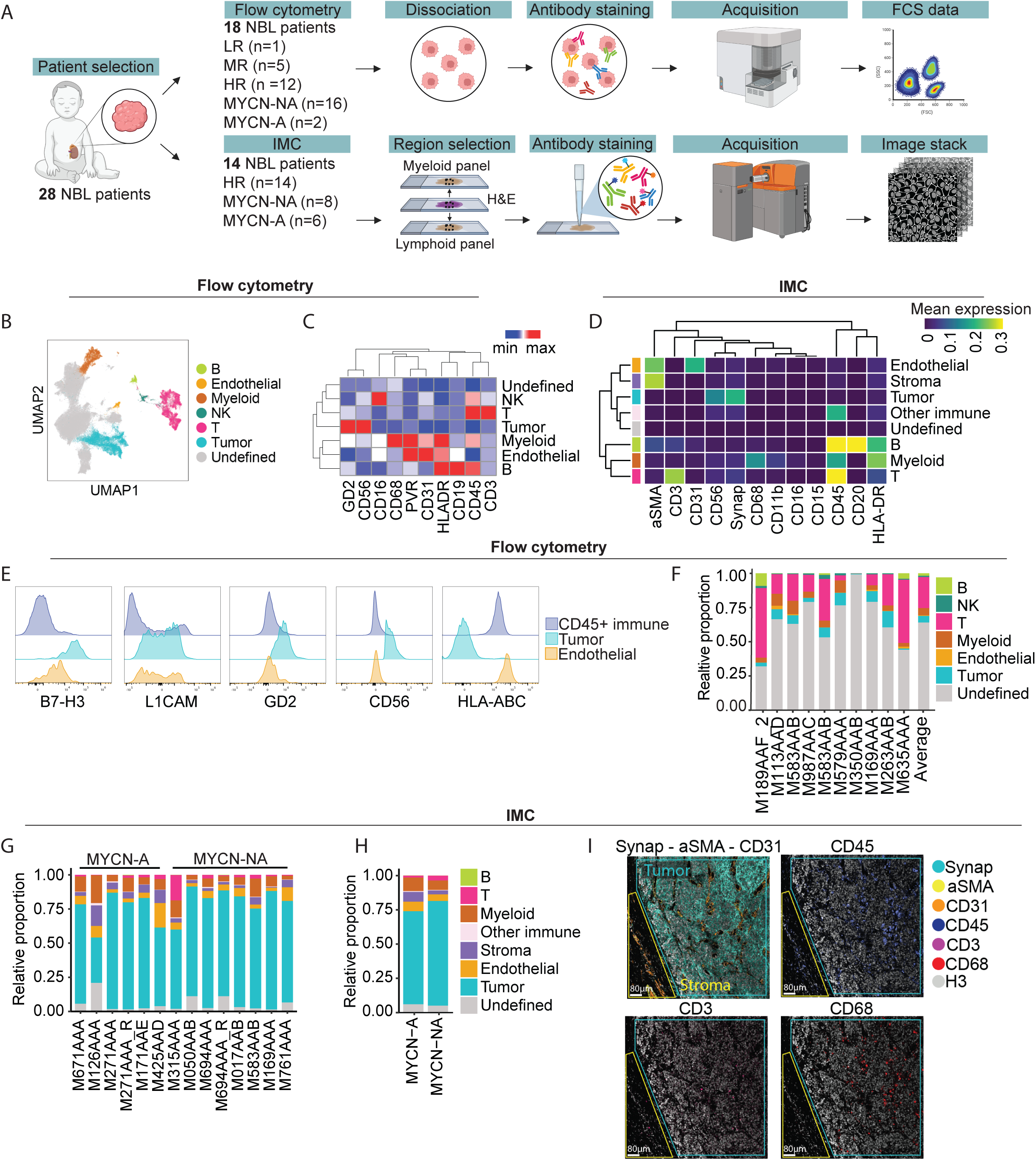
Characterization of the tumor microenvironment of neuroblastoma (A) Experimental design (B) UMAP of main cell types defined by unsupervised clustering of flow cytometry data (C) Heatmap of lineage markers per cell cluster of flow cytometry data (D) Heatmap of mean expression of lineage markers by imaging mass cytometry (IMC) (E) Representative flow cytometry plots with tumor markers (F) Stacked bar plots of relative proportions of cell populations of flow cytometry data, average and per sample (G) Stacked bar plots of relative proportion of cell populations (I) Stacked bar plots of relative proportion of cell populations between MYCN-amplified (MYCN-A) and MYCN non-amplified (MYCN-NA) tumors (propeller differential proportion test, p > 0.05 for all comparisons) (J) Representative IMC images of tumor and stroma/endothelial regions. Synap = synaptophysin (tumor marker), aSMA = alpha-smooth muscle actin (stromal cell marker), CD31 (endothelial marker), CD45 (immune cell marker), CD3 (T cell marker), CD68 (myeloid cell marker)

Based on lineage marker expression, tumor, endothelial, and immune cells were identified across both datasets, including CD3^+^ T cells, (CD19^+^ or CD20⁺) B cells, and HLA-DR^+^/CD68^+^ myeloid cells (**Figures 1B–D, S1 A-B and S2 A-C**). NK cells (CD56⁺^/-^CD16⁺^/-^) were only identified in the flow cytometry dataset, whereas (αSMA^+^) stromal cells were only identified in the IMC dataset. Tumor cells expressed typical neuroblastoma-associated cell surface markers measured by flow cytometry, including B7-H3, L1CAM, GD2, and CD56, while lacking HLA-ABC expression, as previously described^18^ (**Figure 1E**). The major cell populations were consistently identified across patients, with some compositional heterogeneity **(Figure 1F-G and S2 D-G**). Notably, the proportion of tumor cells differed substantially between the flow cytometry (∼5%) and IMC (∼73%) datasets (**Figure 1F-G)**. This likely reflects the differences in tissue sampling strategies, as IMC was performed on tumor-enriched regions of interest (ROIs)^43^, while flow cytometry was performed on dissociated whole-tissue samples with viability gating, thereby excluding tumor cells with reduced viability following chemotherapy^22^ and subject to tumor cell loss during tissue dissociation and thawing of the sample^19^ (**Figure S2B**). The immune composition was broadly comparable between *MYCN*-amplified and *MYCN* non-amplified tumors, with only a trend of lower T cells and higher myeloid and stromal cells in *MYCN* amplified tumors (**Figure 1H**).

Representative images showed exclusive synaptophysin^+^ tumor-rich areas and aSMA^+^/CD31^+^ stromal/vascular-rich areas across the TME (**Figure 1I**). Intriguingly, macrophages (CD68^+^) displayed a dispersed distribution throughout the TME, while T cells (CD3^+^) were less abundant and localized near stromal/vascular tissue. Altogether, neuroblastomas exhibited a heterogeneous TME with spatial organization emerging as an important determinant of immune cell localization.

### Spatial analysis reveals four cellular neighborhoods with distinct immune profiles

To assess the spatial distribution of immune cells in detail, we classified immune cell subsets in the IMC data based on their lineage marker expression. We identified CD4^+^, CD8^+^, and CD4^-^CD8^-^double-negative (DN) T cells, as well as a spectrum of myeloid subsets, comprising monocytes (CD14^+^), macrophages (CD68^+^), CD163^+^ macrophages (CD68^+^CD163^+^), MPO^+^ macrophages (CD68^+^MPO^+^), dendritic cells (CD11c^+^ DCs), neutrophils (CD15^+^CD16^+^MPO^+^), and polymorphonuclear myeloid-derived suppressor cells (PMN-MDSCs; CD15^+^LOX1^+^) (**Figure 2A, S3 A**). Similar populations, including CD68^+^CD163^-^CD206^-^ (DN) and CD163^+/-^CD206^+/-^macrophages, plasmacytoid DCs (pDCs) and conventional DCs (cDCs), were identified by flow cytometry (**Figure S3 B-F**).

**Figure 2.**
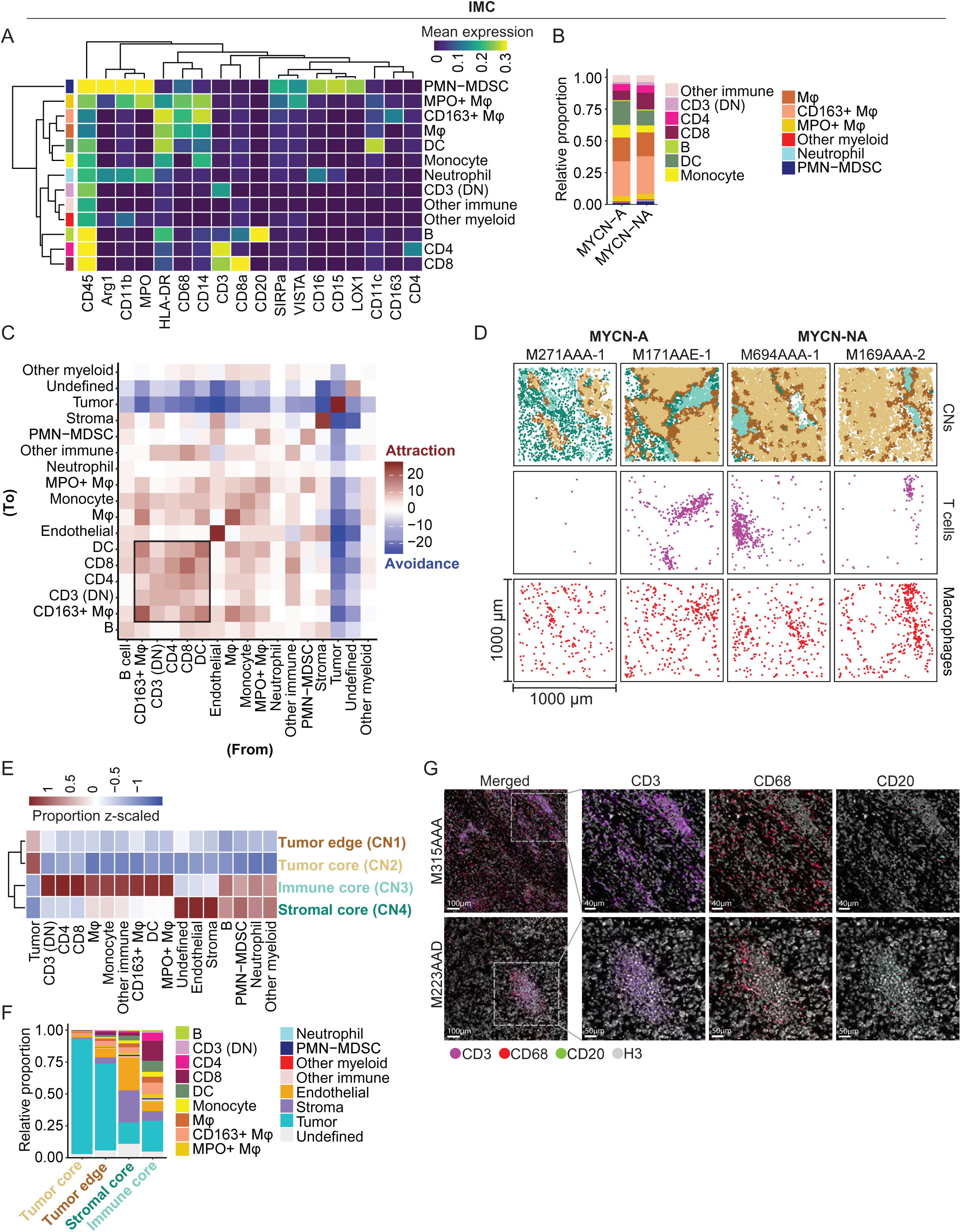
Spatial analysis of cellular neighborhoods in neuroblastoma (A) Heatmap of mean expression of immune lineage markers by imaging mass cytometry (IMC) (B) Stacked bar plots of relative proportion of immune cell populations between MYCN-amplified (MYCN-A) and MYCN non-amplified (MYCN-NA) tumors (propeller differential proportion test, p > 0.05 for all comparisons) (C) Heatmap of the pairwise cell-cell co-colocalization, ranged by attraction (red) versus avoidance (blue) (D) Cellular neighborhood (CN) compositions and spatial distribution of T cells (CD3^DN^, CD4^+^, and CD8^+^) and macrophages (CD163^+^, MPO^+^ and CD68^+^ subtypes) in four representative regions of interest (ROIs) (E) Heatmap visualizing z-scored proportions of cell types across the four cellular neighborhoods annotated in the figure (CN1: tumor edge; CN2: tumor core; CN3: immune core; CN4: stromal core). (F) Stacked barplot visualizing each cellular subset as the proportion of total cells within a specific CN (G) Representative ROIs from immature tertiary lymphoid structure (TLS)-like structures with CD3, CD68, and CD20 staining measured by IMC

All major immune cell populations were detected across patients, with considerable variation in their relative abundances (**Figure S3 G-H**), while overall immune cell composition did not differ significantly between *MYCN*-amplified and non-amplified tumors (**Figure 2B**). The immune compartment was predominantly composed of myeloid cells (**Figure S3 I**), which displayed distinct immunoregulatory profiles. PMN-MDSCs and MPO^+^ macrophages exhibited high expression levels of VISTA and SIRPα (**Figure 2A**), associated with T cell suppression and inhibition of macrophage-mediated phagocytosis^44–46^. Arginase-1 (ARG1) expression, known to suppress T cell function and promote neuroblastoma growth^47,48^, was highest in PMN-MDSCs, followed by neutrophils and MPO^+^ macrophages. CD163^+^ macrophages, generally associated with an immunosuppressive phenotype and poorer outcomes in neuroblastoma^49,50^, also showed SIRPα expression. Thus, PMN-MDSCs, neutrophils, MPO^+^ macrophages, and CD163^+^ macrophages exhibited a broadly immunosuppressive phenotype.

To identify spatial patterns of immune cell infiltration in the TME, we performed co-localization analysis. In line with the observed patterns in **Figure 1I**, co-localization of tumor cells with immune cells was minimal (**Figure 2C, S3 J).** Instead, CD4^+^ and CD8^+^ T cells co-localized with each other, DCs and CD163^+^ macrophages, and to a lesser extent with stromal and endothelial cells. We next asked whether the neuroblastoma TME is organized into recurring, spatially distinct cellular niches. To address this, we computationally defined four cellular neighborhoods (CNs) (**Figure 2D-F, S4 A-B**). The relative abundance of these CNs varied considerably across ROIs and patients, reflecting the substantial spatial heterogeneity of the neuroblastoma TME (**Figure S4 C**). CN1 (tumor edge) and CN2 (tumor core) were dominated by tumor cells, with some CD68^+^ and CD163^+^ macrophages and few T cells (**Figure 2E-F, S4 D**). Macrophages showed a relatively broad distribution across CNs and infiltrated tumor regions more than T cells (**Figure 2E-F, S4 D**). CN4 (stromal core) was enriched in stromal and endothelial cells but also contained both macrophages and T cells (**Figure S4 D**), and showed a trend towards expansion in *MYCN*-amplified tumors (Panel 1: p = 0.24; Panel 2: p = 0.03) (**Figure S4 E**). These stromal niches with ‘entrapped’ immune cells, have previously been associated with high-risk disease, tumor recurrence, and poorer survival^51^.

CN3 (immune core) represented T cell “hotspots” and was characterized by structured aggregates of CD8^+^ and CD4^+^ T cells, with a subset of patients additionally showing scattered B cells within these aggregates (**Figure 2D and G**), suggestive of immature tertiary lymphoid structures (TLS), a pattern previously observed in a subset of metastatic neuroblastoma^52^. Interestingly, one patient (M223AAD), whose tumor was *MYCN* non-amplified, displayed particularly dense TLS-like structures (**Figure 2G**), with a central B cell core surrounded by T cells, with an outer layer of myeloid cells, including dendritic cells, which has previously reported in localized L1 INRG-stage *MYCN* non-amplified neuroblastoma^52^. Altogether, these analyses revealed a spatially structured TME with distinct cellular niches, with immunosuppressive myeloid cells broadly dispersed across niches, and lymphocytes largely confined to tumor edges and immune cores, forming aggregates reminiscent of immature TLS.

### Lymphocytes in neuroblastomas exhibit a tissue-resident effector memory profile with dysfunctional features

To investigate the functional profile of lymphocytes in more detail, we deeply characterized them by spectral flow cytometry. We identified CD8^+^ and conventional CD4^+^ T cells (CD4 conv), CD4^+^FOXP3^+^ regulatory T cells (Tregs), γδT cells, natural killer (NK)T cells, NK cells, and innate lymphoid cells (ILCs) (**Figure 3A-B and S5 A**). Due to the limited number of cells, ILCs were excluded from further analyses. Despite compositional heterogeneity across patients, most subsets were represented in all patients (**Figure 3C, left**). CD8^+^ and CD4^+^ conv cells were the dominant populations, except in patient M451AAD with stage MS neuroblastoma, in whom, remarkably, ∼50% of all lymphocytes consisted of NK cells (**Figure 3C, left**). Comparison of lymphocyte proportions between different sites within the same tumor showed only small differences, suggesting that inter-patient heterogeneity may be more pronounced than intra-patient heterogeneity (**Figure 3C, right**).

**Figure 3.**
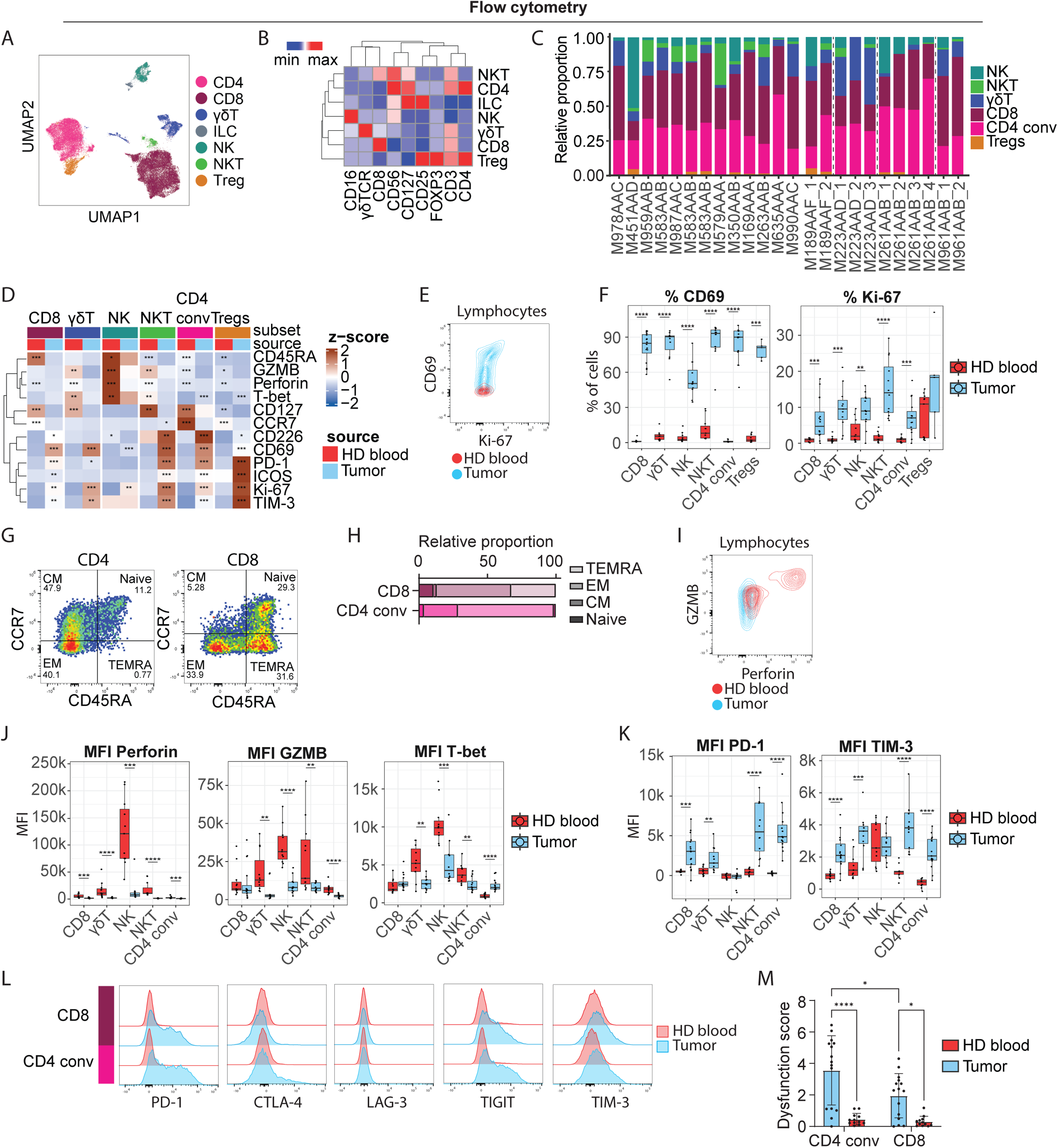
Functional characterization of lymphocytes in neuroblastoma (A) UMAP of T and NK cells defined by unsupervised clustering of flow cytometry data (B) Heatmap of lineage markers of T and NK cells (C) Fraction of T and NK cell populations, left panel one sample per patient, right panel multiple samples per patient (D) Heatmap of all markers significantly different between tumor and blood in 4 or more cell subsets, Pairwise Wilcoxon rank-sum test with BH correction. *p.adj < 0.05, **p.adj < 0.01, ***p.adj < 0.001, ****p.adj < 0.0001 (E) Representative flow cytometry plot of CD69 and Ki-67 expression by lymphocytes in tumor and blood (F) Percentage of CD69 and Ki-67 positive cells per subset, Pairwise Wilcoxon rank-sum test with BH correction. *p.adj < 0.05, **p.adj < 0.01, ***p.adj < 0.001, ****p.adj < 0.0001 (G) Representative flow cytometry plot of four memory phenotypes of CD4^+^ conventional T cells and CD8 T cells, EM = effector memory, TEMRA = effector memory re-expressing CD45RA, CM = central memory (H) Stacked bar plot showing percentage of memory phenotypes of CD4^+^ conventional cells and CD8^+^ T cells (I) Representative flow cytometry plot of granzyme B (GZMB) and perforin of lymphocytes in tumor and blood (J) Mean fluorescent intensity (MFI) of GZMB, perforin, and T-bet on T/NK cells in tumor samples and healthy adult blood samples, Pairwise Wilcoxon rank-sum test with BH correction. *p.adj < 0.05, **p.adj < 0.01, ***p.adj < 0.001, ****p.adj < 0.0001 (K) MFI of PD-1 and TIM-3 on T/NK cells in tumor samples and healthy adult blood samples, Pairwise Wilcoxon rank-sum test with BH correction. *p.adj < 0.05, **p.adj < 0.01, ***p.adj < 0.001, ****p.adj < 0.0001 (L) Representative histograms of PD-1, CTLA-4, LAG-3, TIGIT, and TIM-3 expression by CD8^+^ and CD4^+^ conv T cells in tumor and blood (M) Dysfunction score calculated based on the expression of five inhibitory receptors (PD-1, CTLA-4, LAG-3, TIGIT, TIM-3) of CD8^+^ and CD4^+^ conventional T cells in tumor and blood, 2-way ANOVA with Šídák’s multiple comparisons test; *p<0.05, **p<0.01, ***p<0.001, ****p<0.0001

To decipher the specific functional profile of neuroblastoma-infiltrating T and NK cells, we compared them to reference cells from healthy adult blood. Of note, differences between the adult and pediatric immune systems have been reported^53,54^, which we could not account for in this analysis. Expression of CD69, an early activation marker indicative of tissue residency^55,56^, and Ki-67, a proliferation marker, was higher across tumor-infiltrating lymphocyte subsets than in blood (**Figure 3D-F and S5 B-C**). Expression of CCR7, CD45RA, and CD127, linked to memory phenotypes in T cells, was largely reduced in the tumor (**Figure 3D and S5 B,D-E**). Most CD8^+^ and CD4^+^ conv T cells (54.3% and 70.0%) were CCR7^-^CD45RA^-^ effector memory (EM) cells (**Figure 3G-H**), next to central memory (CM) cells (in CD4^+^ conv T), and terminally differentiated effector memory (TEMRA) cells (in CD8^+^ T), suggesting an advanced differentiation state in CD8^+^ T cells. Together, CD8^+^ and CD4^+^ conv T cells in the tumor displayed a tissue-resident memory phenotype with increased turnover.

Cytotoxicity markers granzyme B, perforin, and T-bet were significantly lower in cytotoxic lymphocyte subsets (CD8^+^, γδT, NK, NKT) in the tumor compared to blood (**Figure 3D, 3I-J and S5 B,D**), indicating a reduction in cytotoxic capacity. Of note, T-bet expression in CD4^+^ conv cells was higher in tumor samples compared to blood, reflecting a Th1 differentiated phenotype (**Figure 3J**). Cytotoxicity of effector cells may be hampered by inhibitory receptor signaling in the TME^57^. Indeed, expression of inhibitory receptors PD-1 and TIM-3 was significantly higher in T cell subsets in the tumor compared to blood (**Figure 3D,K and S5 B-C**), indicative of dysfunctionality. To quantify T cell dysfunctionality across T cell subsets, we used a combined inhibitory receptor expression score (PD-1, CTLA-4, TIM-3, LAG-3, TIGIT)^58^, which proved higher in tumor-infiltrating T cells than blood, and higher in tumor-infiltrating CD4^+^ conv than CD8^+^ T cells (**Figure 3L-M**). Together, this suggests that CD4^+^ conv and CD8^+^ neuroblastoma-infiltrating T cells have distinct dysfunctional features.

Regulatory T cells can also contribute to suppression of effector T cell activity. Although Tregs comprised <2% of the lymphocytes in most patients (**Figure 3C, left**), FOXP3^+^CD25^+^CD127^-^ Tregs in tumor samples showed higher expression of their signature transcription factor FOXP3, of activation markers ICOS and CD137, and of inhibitory receptors CTLA-4 and TIGIT compared to Tregs from the blood (**Figure S5 F-I**). CTLA-4 and TIGIT are part of the core signature of Tregs, and enhanced expression is associated with enhanced suppressive capacity^59–61^, hinting towards an activated effector Treg phenotype in the tumor. Overall, these data indicate that tumor-infiltrating T cells exhibit a tissue-resident memory profile with subset-specific dysfunctional features, including activated Tregs.

### The spatial niche and local tumor cell density define CD8^+^ T cell functional states

To assess whether the observed (dys)functional properties of T cells varied by spatial niche and validate tissue residency of T cells, we investigated the expression of markers representing proliferation (Ki-67), stemness/exhaustion (TCF1/7, TOX)^62–65^, and cytotoxic capacity (T-bet, granzyme B) across the four defined CNs in our IMC data.

Ki-67 expression was highest in the immune core, indicative of local proliferation, with higher expression in CD4^+^ than CD8+ T cells (**Figure 4A-B, S6 A**). Concordantly, TCF1/7 expression, associated with stemness, self-renewal capacity, and progenitor exhaustion when co-expressed with TOX ^62–65^, decreased gradually from the immune core towards tumor-rich CNs (tumor core and tumor edge), consistent with a recent study showing enrichment of TCF7^+^, stem-like T cells in immune-rich niches in high-risk neuroblastoma^66^ (**Figure 4A, S6 A**). Although TOX expression was low ov, it was highest in CD4^+^ T cells in the immune core, suggestive of local progenitor exhaustion (TCF1/7^+^TOX^+^)^62–64^, possibly driven by local repetitive antigen stimulation. Interestingly, despite overall low granzyme B expression in tumor-infiltrating CD8^+^ T cells compared to healthy blood (**Figure 3J**), CD8^+^ T cells in the tumor core showed an enhanced cytotoxic profile compared to other CNs, marked by granzyme B expression (**Figure 4A-B, S6 A**), suggestive of cytotoxic activation upon tumor contact.

**Figure 4.**
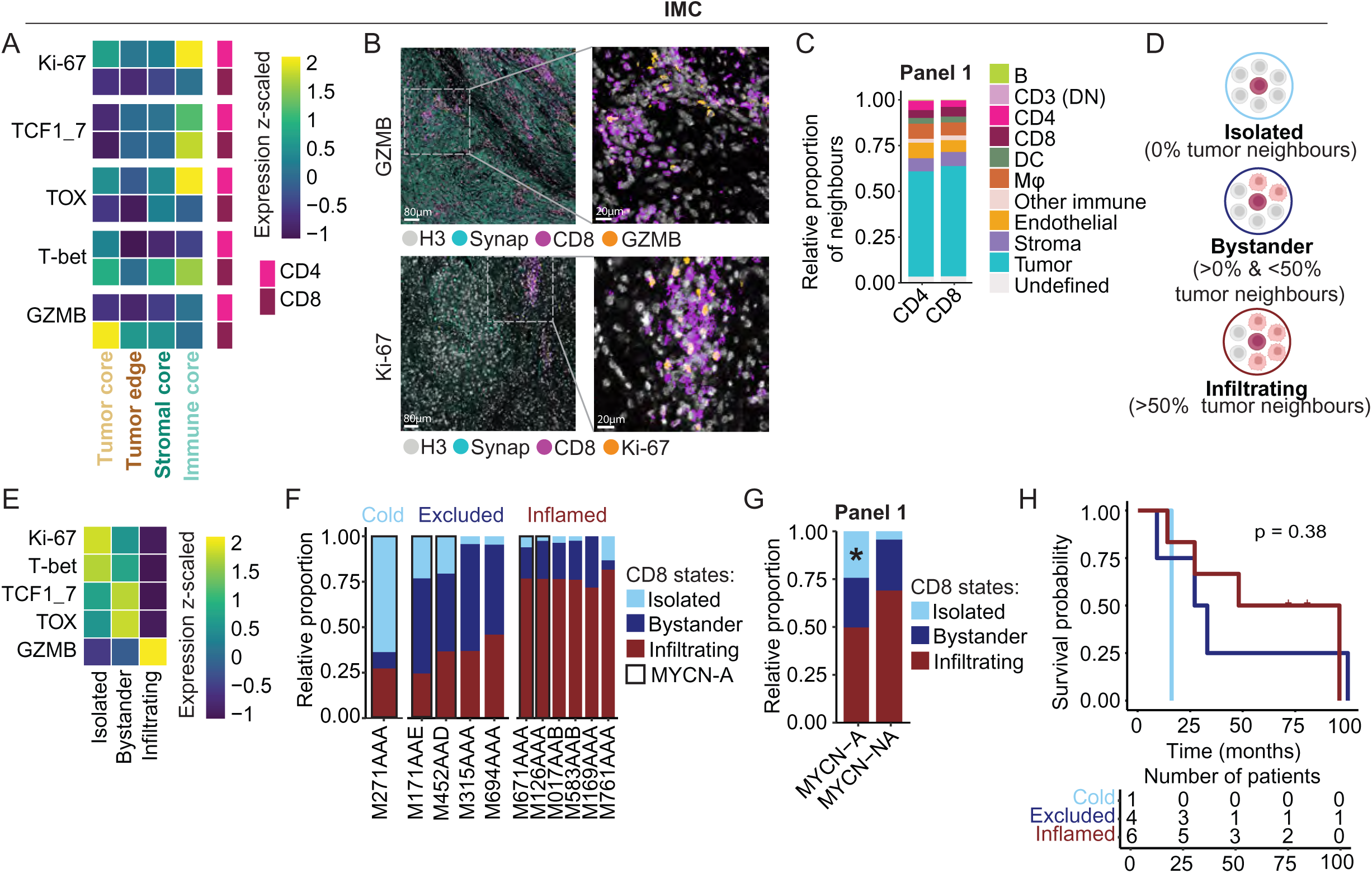
T cell functional states defined by spatial niches and tumor cell density (A) Heatmap of z-scaled expression (scaled per marker, across CD4^+^ and CD8^+^ T cells and CNs) of Ki-67, TCF1/7, TOX, T-bet, and granzyme B (GZMB) in CD8^+^ and CD4^+^ T cells across the four cellular neighborhoods measured by imaging mass cytometry (IMC) (B) Representative regions of interest (ROIs) showing Ki-67 and GZMB staining measured by IMC (C) Relative proportion of CD4^+^ and CD8^+^ T cell neighbours in panel 1 of IMC data (propeller differential proportion test, p > 0.05 for all comparisons) (D) Schematic illustrating the classification of CD8 T cells into spatial states based on the proportion of neighbouring tumor cells (created with BioRender.com) (E) Mean expression of Ki-67, TCF1/7, T-bet, TOX, and GZMB per CD8 T cell state (F) Stacked bar plots of the relative proportion of CD8 T cell states, and stratification of patients into cold, excluded or inflamed tumors based on the highest abundant CD8 T cell state (G) Stacked bar plots of the relative proportion of CD8 T cell states between MYCN-amplified (MYCN-A) and MYCN non-amplified (MYCN-NA) tumors (*<p,0.05; propeller differential proportion test) (H) Kaplan-Meier curves of overall survival (OS) for patients into tumor groups (cold, excluded, inflamed); p values were estimated using a log rank test

Given that CD8^+^ T cell cytotoxic activity may be linked to tumor proximity, and that >60% of CD8+ T cells neighbors were tumor cells (**Figure 4C, S6 B**), we next examined the extent of direct tumor contact and its effect on the CD8^+^ T cell phenotype. We defined three spatial states of CD8^+^ T cells according to the proportion of neighboring tumor cells, adapted from Meyer et al.^42^: ‘isolated’ (0%), ‘bystander’ (0-50%), and ‘infiltrating’ (>50%) (**Figure 4D**). While infiltrating CD8^+^ T cells were surrounded by tumor neighbors, isolated and bystander CD8^+^ T cells were predominantly neighbored by other T cells (**Figure S6 C-D**). Phenotypically, infiltrating CD8^+^ T cells showed the highest granzyme B expression, consistent with cytotoxic activity, whereas bystander CD8^+^ T cells displayed a more progenitor-exhausted profile (TCF1/7^+^TOX^+^), and isolated CD8^+^ T cells, lacking tumor contact, were the most proliferative (**Figure 4E**). Thus, CD8^+^ T cell differentiation shows a spatial gradient, with stem-like, self-renewing CD8^+^ T cells enriched in the immune core, a more differentiated/exhausted-associated phenotype at the tumor-immune interface, and a cytotoxic phenotype among tumor-infiltrated CD8^+^ T cells.

In addition to these phenotypic differences, the spatial context of CD8^+^ T cells has been linked to clinical outcome, with CD8^+^ T cell infiltration and localization showing prognostic relevance in neuroblastoma^10,67^ and tumor density around CD8^+^ T cells showing prognostic associations in other tumors^42^. We therefore classified patients by their dominant CD8^+^ T cell state (**Figure 4F**). Remarkably, *MYCN*-amplified tumors had more isolated CD8^+^ T cells (panel 1: p=0.029; panel 2: p=0.064), and *MYCN* non-amplified tumors showed a trend of more infiltrating CD8^+^ T cells (**Figure 4G and S6 E**), confirming previously observed T cell exclusion in *MYCN*-amplified tumors^9,12^. Moreover, patients with inflamed tumors showed a trend of better overall survival (OS), as also observed in other tumor types^68–70^ (**Figure 4H and S6 F**). Altogether, spatial localization within defined cellular niches and local tumor cell density appear to be closely linked to T cell functional states, with tumor-engaged CD8^+^ T cells exhibiting a more cytotoxic phenotype.

### CD163^+^ macrophages are the dominant immunosuppressive interaction partners of T cells

Since the distinct cellular neighborhoods may not only dictate T cell functional states, but also their cellular interactions, we examined significant T cell-TME interactions occurring in at least 20% of ROIs across the four CNs (**Figure 5A and S7A**). Within the tumor and stromal cores, recurrent T cell interactions were largely absent, consistent with the limited T cell infiltration observed in this niche. At the tumor edge, CD8^+^ T cells had significant interactions with endothelial, tumor, and stromal cells, as well as CD163^+^ macrophages. The immune core displayed the richest interaction profile, characterized by abundant T cell-T cell interactions next to CD8^+^ T cells interactions with CD163^+^ macrophages and DCs. This suggests that T cells are subject to distinct cellular interactions within each CN, including potentially inhibitory interactions with immunosuppressive myeloid cells.

**Figure 5.**
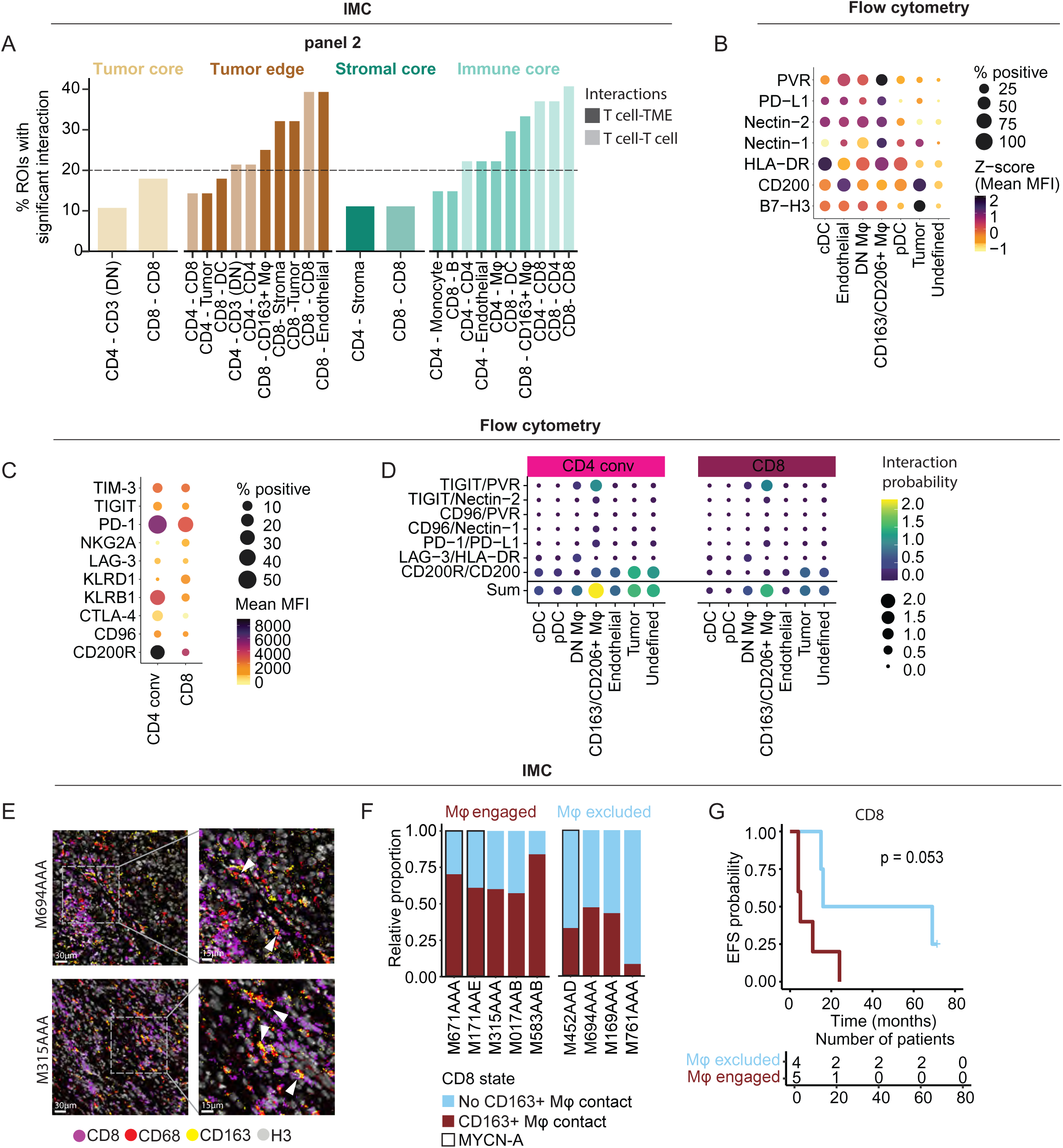
Immunosuppressive interactions between CD163^+^ macrophages and T cells in the neuroblastoma tumor microenvironment (A) Pairwise CD8^+^ and CD4^+^ T cell interactions were calculated for each cellular neighborhood. Bar plots show the 10 most frequent significant CD8^+^ and CD4^+^ T cell interaction occurring in more than 10% of regions of interest (ROIs) (B) Inhibitory ligand expression (percentage of marker expressing cells and z-scored mean fluorescent intensity (MFI)) by myeloid cells, tumor cells, endothelial cells, and undefined cells measured by flow cytometry (C) Inhibitory receptor expression (percentage of marker expressing cells and MFI) by T/NK cells measured by flow cytometry (D) Bubble plot of interaction probability of inhibitory receptor/ligand pairs, calculated as described in detail in the methods section, between CD4^+^ conventional and CD8^+^ T cells and myeloid cells, endothelial cells, tumor cells, and undefined cells, based on flow cytometry data (E) Representative ROI images depicting CD8^+^ T cell interaction with CD163^+^ macrophages within the tumor microenvironment (F) Stacked bar plot showing relative proportions of CD8 T cells states based on their contact with CD163^+^ macrophages, stratifying patients into macrophage engaged or excluded based on the predominant phenotype. (G) Kaplan Meier curve showing the event free survival (EFS) probability for patients that are stratified into macrophage engaged or excluded based on their CD8 state; p values were estimated using a log rank test

To interrogate which known inhibitory ligand-receptor interactions T cells are subject to in the neuroblastoma TME, we analyzed their detailed protein expression by flow cytometry. Tumor cells expressed high levels of B7-H3 (receptor yet unknown) and CD200R ligand CD200 (**Figure 5B**). Compared to other cells in the TME, particularly CD163^+^/CD206^+^ macrophages expressed high levels of CD96 ligand Nectin-1, TIGIT ligands Nectin-2 and PVR, and PD-1 ligand PD-L1 (**Figure 5B**). CD4^+^ conv T cells expressed high KLRB1, CD200R, and PD-1 (**Figure 5C**), while in concordance with our previous observation, CD8^+^ T cells had a lower expression of these receptors (**Figure 3L-M and figure 5C**).

To model the interaction probability between these inhibitory ligands and their receptors, we calculated – for each receptor-ligand pair and for each combination of interacting populations – an interaction probability score based on the MFI, the population size, and the percentage of ligand/receptor-expressing cells. Distinct T cell subsets showed similar interaction patterns (**Figure 5D and S7 B**), with the strongest predicted interactions being the CD200/CD200R and TIGIT/PVR axes (**Figure 5D and S7 B**). Overall, CD163^+^/CD206^+^ macrophages had the highest sum of inhibitory interactions with T cells (**Figure 5D**), indicating that they are likely the dominant inhibitory interaction partners for T cells in the TME. In line with this, distinct clusters of CD163^+^ macrophages co-localizing with CD8^+^ T cells were observed in the IMC data (**Figure 5E**).

To quantify these interactions and their prognostic relevance, we classified CD8^+^ T cells as having either ‘contact’ or ‘no contact’ with CD163^+^ macrophage neighbors and assessed whether this classification was associated with patient outcomes (**Figure 5F**). Patients with predominantly CD163^+^ macrophage-engaged CD8^+^ T cells showed a trend towards worse event free survival (EFS) and OS (**Figure 5F-G and S7C**). A similar prognostic pattern was observed for CD4^+^ T cells (**Figure S7 D-F**). Collectively, these findings identify CD163⁺ macrophages as the dominant immunosuppressive interaction partners of T cells, with increased macrophage–T-cell engagement being associated with poorer patient survival.

### Nectin-2 expression on macrophages within T cell-rich niches is associated with poor **patient outcomes**

Extending the association between CD163⁺ macrophage-T cell interactions and patient survival, we next asked whether phenotypic differences within the CD163⁺ macrophage compartment also carried prognostic significance. In the flow cytometry data, among these phenotypic markers, CD163 expression itself by CD163^+^/CD206^+^ macrophages significantly correlated with worse EFS (R =-0.93, p = 0.00025) (**S7 G-H**). Similarly, among the inhibitory ligands, Nectin-2 expression correlated negatively with survival (R =-0.78, p = 0.017) (**Figure 6A-C**). Nectin-2, expressed on CD163^+^ macrophages, binds the inhibitory receptor TIGIT on T cells, thereby contributing to T cell suppression, as we previously reported^22^.

**Figure 6.**
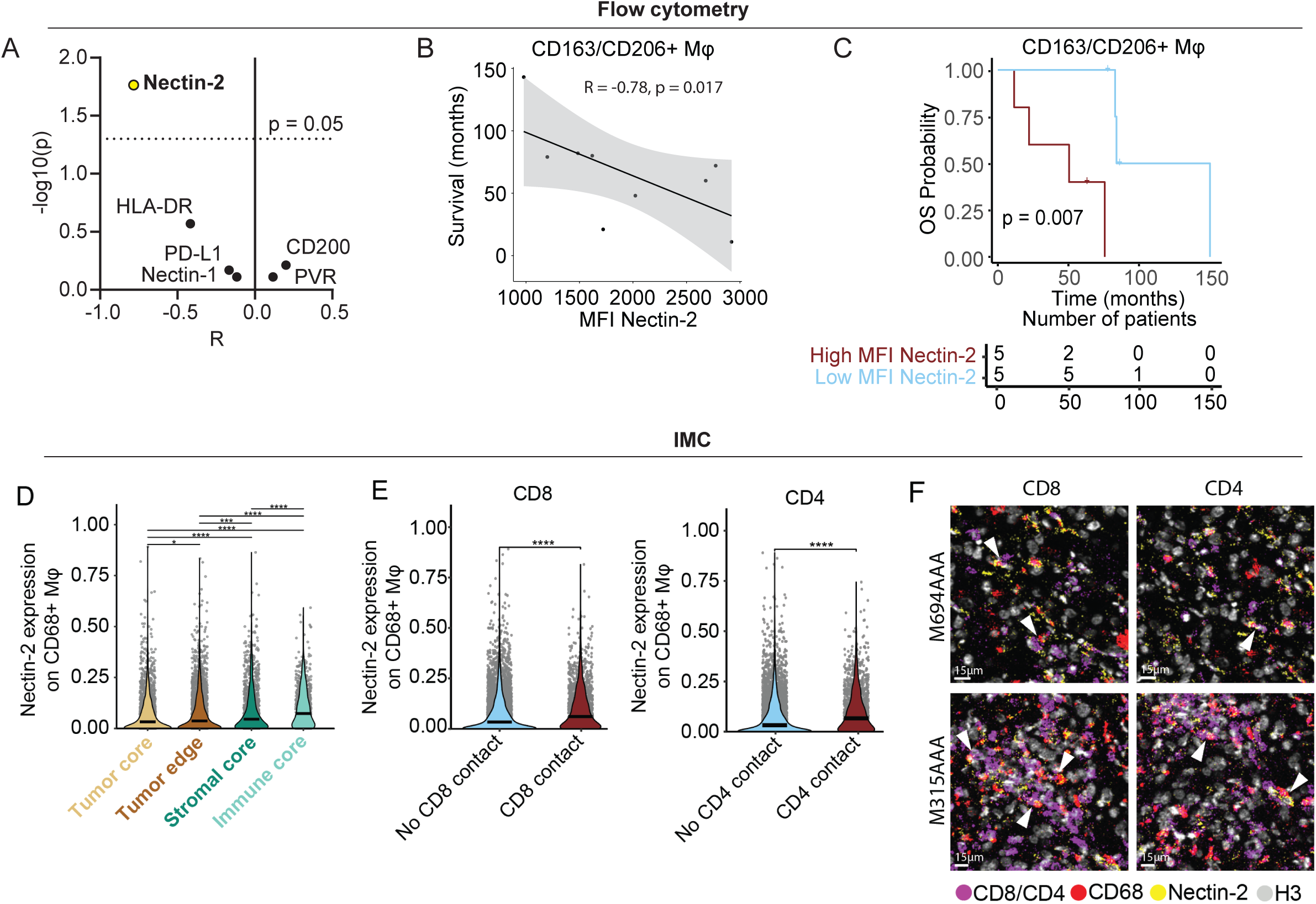
Nectin-2-expressing macrophages associated with patient survival (A) Kaplan Meier curve showing the event free survival (EFS) probability for patients with high and low MFI CD163 on CD163^+^/CD206^+^ macrophages measured by flow cytometry; p values were estimated using a log rank test (B) Spearman correlations between patient survival and expression of inhibitory ligands on CD163^+^/CD206^+^ macrophages, showing the correlation coefficient (R) and-log10(p-value), based on flow cytometry data (C) Spearman correlation between patient survival and Nectin-2 expression on CD163+/CD206+ macrophages, based on flow cytometry data (D) Kaplan Meier curve showing the overall survival (OS) probability for patients with high and low MFI Nectin-2 on CD163^+^/CD206^+^ macrophages measured by flow cytometry; p values were estimated using a log rank test (E) Nectin-2 expression on CD68^+^ macrophages across the four cellular neighborhoods (CNs). Statistical significance between CNs was assessed using pairwise Wilcoxon rank-sum test with BH correction. *p.adj < 0.05, **p.adj < 0.01, ***p.adj < 0.001, ****p.adj < 0.0001 (F) Nectin-2 expression on CD68^+^ macrophages in contact with versus no contact with CD8+ or CD4+ T cells. Groups were compared using Wilcoxon rank-sum test. *p.adj < 0.05, **p.adj < 0.01, ***p.adj < 0.001, ****p.adj < 0.0001 (G) Representative images of ROIs depicting CD8^+^ or CD4^+^ T cells in proximity to Nectin-2 expressing CD68^+^ macrophages

To decipher where this immunosuppressive interaction is most likely to occur, we next examined Nectin-2 expression on macrophages across the four defined CNs in the IMC data. Although a small difference, Nectin-2 expression was significantly higher in the immune core than in the tumor core (median normalized expression: 0.0731 vs 0.0324, 2.3-fold change; padj = 1.38e-23) (**Figure 6D**). Given that the immune core is rich in T cells, we next asked whether this elevated Nectin-2 expression on macrophages depends on direct T cell interaction. Indeed, macrophages with direct T cell contact showed significantly higher Nectin-2 expression than those without T cell contact (CD8 contact: median normalized expression: 0.0612 vs 0.0341, 1.8-fold change; padj = 5.01e-28; CD4 contact: median normalized expression: 0.0676 vs 0.0332, 2.0-fold change, padj = 2.08e-36) (**Figure 6E-F**). Collectively, these results reveal that Nectin-2 expression on immunosuppressive macrophages is enriched within macrophage-T cell-rich niches, increasing the likelihood of interacting with TIGIT^+^ T cells *in situ,* and correlating with worse clinical outcomes. Thus, the TIGIT/Nectin-2 interaction emerges as a defining feature of the neuroblastoma TME and a compelling therapeutic target.

## DISCUSSION

Despite substantial progress in improving immunotherapy for neuroblastoma, to date, the clinical benefit of T cell-based therapies remains limited, especially in patients with a high disease burden^2,13,14^. Increasing evidence suggests that the immunosuppressive TME plays a key role in treatment failure^23^. To gain a deeper understanding of the spatial context of cellular interactions in the neuroblastoma TME, we combined imaging mass cytometry with high-dimensional spectral flow cytometry data. We identified a spatially structured TME characterized by the exclusion of T cells from tumor-dominant regions. Instead, T cells were retained in immature TLS-like immune aggregates embedded within the tumor and stromal cores. There, CD8^+^ T cells in particular were subject to pronounced immunoregulatory interactions with CD163^+^ macrophages, including the TIGIT/Nectin-2 axis, which was predictive of patient outcomes.

While neuroblastoma is typically regarded as an immunologically cold tumor compared to many adult cancers^71^, we observed T cells in the majority of patients, both in this study and in a previous systematic meta-analysis of T cell infiltration across pediatric tumors^71^. The presence of T cells in this study on post-treatment samples may be related to prior chemotherapy, as cytotoxic agents have been shown to support immunogenic cell death, and thereby T cell attraction to tumor sites^43,72^. Overall, however, T cells were largely excluded from tumor-dominant niches and instead confined to immune aggregates, localized within the tumor and stromal cores. The stromal core (CN4) was more prevalent in *MYCN*-amplified tumors, consistent with a previously described “stroma-secluded immune niche” linked to poorer OS and potential resistance to conventional therapies in high-risk neuroblastoma^51^. Interestingly, in a subset of patients, the T cell-rich immune aggregates displayed a spatial pattern of immature TLS-like structures, characterized by disorganized B and T cell aggregates. These findings are similar to what has been previously described in localized *MYCN* non-amplified neuroblastoma, which was linked to more effective antitumor responses^52^. Together, these spatial patterns underscore the heterogeneity of immune organization in neuroblastoma and its potential relevance for clinical outcomes.

The observed immune aggregates were primarily characterized by the presence of CD4^+^/CD8^+^ T cells, B cells, macrophages, and DCs, and high expression of tissue residency and effector phenotype markers in T cells. However, effector functions of T cells remained impaired, reflected by reduced perforin and granzyme B expression in CD8^+^ T cells, as well as increased expression of inhibitory receptors (TIM-3 and PD-1) mainly in CD4^+^ T cells, consistent with an exhausted/dysfunctional phenotype and in line with previous neuroblastoma studies^20,22^. The immune aggregates likely support effective (and possibly repetitive) antigen presentation for CD4^+^ T cells, in line with the observed CD4^+^-myeloid cell interactions, and the exhausted/dysfunctional profile observed in CD4^+^ T cells. On the other hand, spatial exclusion of CD8^+^ T cells from tumor cores, and thus lack of antigen stimulation, may explain the lack of inhibitory receptor expression. The few ‘inflamed’ neuroblastomas with CD8^+^ T cells infiltrating tumor cores showed a trend towards improved OS, whereas T cell-excluded tumors were associated with poorer OS, consistent with a previously described T cell-inflamed signature associated with improved OS in high-risk neuroblastoma^67^. Notably, although less abundant, these tumor-infiltrating CD8^+^ T cells displayed a more cytotoxic profile, marked by higher levels of granzyme B expression, potentially contributing to the improved survival in these patients with ‘inflamed’ tumors.Although these findings are based on a limited number of patients and require further validation in larger cohorts, they suggest that T cell localization may be clinically relevant.

CD163^+^ macrophages were identified as the main immunosuppressive interaction partner for CD8^+^ T cells, with this interaction significantly predicted within the tumor edge and immune core. Ligand–receptor analysis suggested a high interaction probability between CD8^+^ T cells and these immunosuppressive CD163^+^ macrophages via the TIGIT/PVR and TIGIT/Nectin-2 axis, confirming previous studies^22,73^. Notably, CD8^+^ T cells within the tumor edge and immune core displayed a more exhausted phenotype, potentially driven by sustained CD163^+^ macrophage interaction mediated through this TIGIT/PVR/Nectin-2 axis, consistent with the same TAM-CD8^+^ T cell TIGIT-Nectin-2 axis linked to exhaustion in an adult solid tumor^74^. Next to this, both CD163 and Nectin-2 expression on macrophages were independently associated with worse survival, further substantiating the clinical relevance of this immunosuppressive interaction.

A unique strength of our study is the comprehensive characterization of the spatial context of the neuroblastoma immune TME, at the single-cell and protein levels, by combining high-dimensional spectral flow cytometry with IMC, enabling both in-depth phenotypic profiling and spatial mapping of immune populations. Nevertheless, several methodological considerations should be taken into account when interpreting our findings. For the spectral flow cytometry analyses, we compared effector functions of lymphocytes in neuroblastomas with those from healthy adult blood, while differences between adult and pediatric blood have been described, as T and NK cells from children are still developing^53,54^. Therefore, comparisons between cells from tumor samples and blood samples should be interpreted with caution. In addition, tissue dissociation potentially modified cell compositions, induced stress responses, and does not allow to distinguish tissue-infiltrated from circulating immune cells^19^. For IMC, our findings underscore the importance of ROI selection, as the neuroblastoma TME displayed substantial spatial heterogeneity. Local immune niches may therefore be overlooked when only limited tissue regions are assessed. Additionally, the relatively small cohort sizes and inter-patient heterogeneity in the TME limit generalizable conclusions regarding prognostic associations.

Altogether, our study highlights several opportunities for therapeutic intervention to enhance the efficacy of T cell-based immunotherapies. First, strategies to enhance T cell infiltration into tumor cores are a promising avenue. Since the extracellular matrix (ECM) constitutes a major physical barrier to T cell infiltration into stroma-rich solid tumors, several ECM-degrading engineering approaches have been developed. CAR-T cells engineered to express heparanase, which degrades heparan sulfate proteoglycans in the ECM, showed improved tumor infiltration and survival of tumor models, including neuroblastoma xenografts^75,76^, and ablation of DDR1, a collagen receptor, enhanced T cell infiltration into the tumor core in a triple-negative breast cancer model^77^. Beyond enzymatic ECM degradation, synthetic “velocity receptors” that bind self-produced cytokines increase the migratory phenotype of CAR-T cells, thereby boosting tumor infiltration and survival in solid tumor models^78^. FAP-targeted CAR-T cells or antibodies can similarly deplete or reprogram FAP^+^ stromal cells to reshape the stromal architecture and enhance immune infiltration^79,80^. Second, targeting the immunosuppressive CD163^+^ macrophage compartment represents another therapeutic opportunity. CSF1R inhibition is the most clinically advanced strategy to deplete or repolarize tumor-associated macrophages (TAMs), given that CSF1 is essential for macrophage survival and differentiation. CSF1R blockade (RG7155/emactuzumab) reduced CSF1R^+^CD163^+^ TAMs and shifted the CD8/CD4 T cell ratio, with objective responses in a phase I trial, and in a neuroblastoma xenograft model, anti-CSF1R antibody depletion of macrophages enhanced chemotherapy efficacy^81,82^. Alternatively, CD163^+^ TAMs can be selectively targeted using CD163-directed nanoparticles delivering doxorubicin, which inhibited tumor growth and was associated with a more immunostimulatory TME^83,84^. More recently, TREM2, a receptor expressed on a subset of TAMs, has emerged as a target. TREM2-targeting, IL-12-expressing CAR-T cells depleted TAMs and remodeled the TME, including enhanced T cell infiltration, in solid tumor models^85,86^. In addition, tumor immune cell targeting chimeras (TICTACs), which are bifunctional small molecule–antibody conjugates that selectively bind TAMs, were developed to deplete immune checkpoint receptors such as SIRPα on TAMs, thereby increasing macrophage-dependent tumor killing in vivo^87^. Third, checkpoint inhibition, including but not limited to TIGIT blockade, offers additional opportunities to restore T cell function. In neuroblastoma, combined TIGIT/PD-L1 blockade significantly enhanced anti-GD2 efficacy *in vivo*^22,88^. In adult cancers, TIGIT-based chimeric switch receptors, which convert an inhibitory TIGIT signal into a costimulatory one, have been engineered to improve T cell antitumor function^89^, and TIGIT blockade is being explored to overcome T cell exhaustion following BCMA-CAR-T therapy in multiple myeloma^90^. TIGIT base-editing was also recently shown to enhance the function of low-avidity engineered T cells in pancreatic ductal adenocarcinoma^91^. More broadly, combination immune checkpoint blockade has demonstrated superior efficacy over single-agent blockade across multiple adult tumor types^92^, and combining checkpoint blockade with genetically engineered T cell therapies, for example, CAR-T cells secreting PD-1-blocking scFvs^93^, is increasingly being explored.

In summary, this study provides a comprehensive phenotypic and spatial characterization of cellular immune interactions in the neuroblastoma TME, revealing T cell exclusion and strong inhibitory CD163^+^ macrophage - CD8^+^ T cell interactions in tumor edges and immune niches, including the Nectin-2/TIGIT axis, which likely contribute to CD8^+^ T cell dysfunction. These findings provide a rationale for combinatorial immunotherapeutic strategies that enhance T cell infiltration into tumor cores, reprogram or eradicate the suppressive myeloid compartment and restore T cell effector function by targeting immunosuppressive checkpoints such as TIGIT/Nectin-2.

## FIGURE LEGENDS

**Suppl Figure 1**

(A) Schematic representation of the cell lineage assignment strategy used for imaging mass cytometry (IMC)

(B) Representative IMC images of all markers per panel

**Suppl Figure 2**

(A) UMAP with sample distribution of flow cytometry data

(B) Flow cytometry gating strategy of main cell types

(C) Heatmap of mean expression of lineage markers of panel 1 by imaging mass cytometry (IMC)

(D) Stacked bar plot showing numbers of cells per sample of flow cytometry data

(E) Stacked bar plots of relative proportion of cell populations within panel 1 of the IMC data

(F) Stacked bar plots of absolute cell population counts within panel 2 of the IMC data

(G) Stacked bar plots of absolute cell population counts within panel 1 of the IMC

**Suppl Figure 3**

(A) Heatmap of mean expression of immune markers panel 1 of imaging mass cytometry (IMC) data

(B) UMAP of myeloid cells defined by unsupervised clustering of flow cytometry data

(C) Heatmap of lineage markers of myeloid cells in flow cytometry data

(D) Flow cytometry gating strategy of myeloid cells

(E) Fraction of myeloid populations per patient in flow cytometry data

(F) Stacked bar plot of number of cells of myeloid populations in flow cytometry data

(G) Stacked bar plots of relative proportion of immune populations within panel 2 of the IMC data

(H) Stacked bar plots of relative proportion of immune populations within panel 1 of the IMC data

(I) Stacked bar plots of relative proportion of lymphoid-to-myeloid cell ratio between in the two IMC panels (propeller differential proportion test, p > 0.05 for all comparisons)

(J) Heatmap of significant pairwise cell-cell attraction (red) or avoidance (blue) across all cell types in panel 1 of the IMC data

**Suppl Figure 4**

(A) Heatmap visualizing z-scored proportions of cell types within each of the four cellular neighborhoods annotated in the figure (CN1:tumor edge; CN2: tumor core; CN3: immune core; CN4: stromal core) in panel 1 of the imaging mass cytometry (IMC) data

(B) Stacked bar plot visualizing each cellular subset as the proportion of total cells within a specific CN in panel 1

(C) Stacked bar plots of relative proportion of cellular neighborhoods across regions of interest (ROIs) between MYCN-amplified (MYCN-A) and MYCN non-amplified (MYCN-NA) tumors within both IMC panels

(D) Boxplots comparing cell type frequencies between cellular neighborhoods. Data represent, for each cellular subset, the percentage of total cells within that CN, as an average value per patient. Each patient is represented as one data point. Pairwise Wilcoxon rank-sum test with BH correction. *p.adj < 0.05, **p.adj < 0.01, ***p.adj < 0.001, ****p.adj < 0.0001

(E) Stacked bar plots of relative proportion of cellular neighborhoods between MYCN-amplified (MYCN-A) and MYCN non-amplified (MYCN-NA) tumors in both panels of the IMC data (*p<0.05; propeller differential proportion test)

**Suppl Figure 5**

(A) Flow cytometry gating strategy of T/NK cells

(B) Volcano plots of all T and NK cells showing differently expressed markers between tumor and healthy adult blood, Mann-Whitney U test; adjusted p<0.05

(C) UpsetR plot of markers that are significantly higher expressed in the tumor (adjusted p<0.05), across the various lymphocyte populations

(D) UpsetR plot markers that are significantly higher expressed in blood (adjusted p<0.05), across the various lymphocyte populations

(E) Mean fluorescent intensity (MFI) of CCR7, CD45RA, and CD127 on T/NK cells in tumor samples and healthy adult blood samples, Pairwise Wilcoxon rank-sum test with BH correction. *p.adj < 0.05, **p.adj < 0.01, ***p.adj < 0.001, ****p.adj < 0.0001

(F) Heatmap of all markers significantly different between tumor and blood in 3 or less cell subsets, Pairwise Wilcoxon rank-sum test with BH correction. *p.adj < 0.05, **p.adj < 0.01, ***p.adj < 0.001, ****p.adj < 0.0001

(G) MFI of FOXP3 in Tregs in tumor samples and healthy adult blood samples, Mann-Whitney U test

(H) MFI of CD137, CTLA-4, and TIGIT on Tregs and CD4^+^ conv cells in tumor samples and healthy adult blood samples, Pairwise Wilcoxon rank-sum test with BH correction. *p.adj < 0.05, **p.adj < 0.01, ***p.adj < 0.001, ****p.adj < 0.0001

(I) MFI of ICOS on Tregs and CD4^+^ conv cells in tumor samples and healthy adult blood samples, Pairwise Wilcoxon rank-sum test with BH correction. *p.adj < 0.05, **p.adj < 0.01, ***p.adj < 0.001, ****p.adj < 0.0001

**Suppl figure 6**

(A) Heatmap of mean expression of Ki-67, TCF1/7, TOX, T-bet, and granzyme B (GZMB) in CD8^+^ and CD4^+^ T cells measured by imaging mass cytometry (IMC)

(B) Relative proportion of cellular neighbours of CD4^+^ and CD8^+^ T cells in panel 2 of imaging mass cytometry (IMC) data (propeller differential proportion test, p > 0.05 for all comparisons)

(C) Relative proportion of bystander and isolated CD8 T cell neighbours in panel 1 of IMC data

(D) Relative proportion of bystander and isolated CD8 T cell neighbours in panel 2 of IMC data

(E) Stacked bar plots of the relative proportion of CD8 T cell states between MYCN-amplified (MYCN-A) and MYCN non-amplified (MYCN-NA) tumors in panel 2 (propeller differential proportion test, p > 0.05 for all comparisons)

(F) Kaplan-Meier curves of OS for patients into tumor groups (cold, excluded, inflamed) in panel 1; p values were estimated using a log rank test

**Suppl Figure 7**

(A) Pairwise CD8^+^ and CD4^+^ T cell interactions were calculated for each cellular neighborhood (CN1:tumor edge; CN2: tumor core; CN3: immune core; CN4: stromal core). Bar plots show the 10 most frequent significant CD8^+^ and CD4^+^ T cell interaction occurring in more than 10% of ROIs within panel 1 of the imaging mass cytometry (IMC) data

(B) Bubble plot of interaction probability of inhibitory receptor/ligand pairs between γδT cells, NK cells or NKT cells and myeloid cells, endothelial cells, tumor cells, or undefined cells measured by flow cytometry

(C) Kaplan Meier curve showing the overall survival (OS) probability for patient that are stratified into macrophage engaged or excluded based on their CD8 state

(D) Stacked bar plot showing relative proportions of CD4^+^ T cells states based on their contact with CD163^+^ macrophages, stratifying patients into macrophage engaged or excluded based on the predominant phenotype

(E) Kaplan Meier curve showing the EFS probability for patient that are stratified into macrophage engaged or excluded from based on their CD4^+^ T cell state

(F) Kaplan Meier curve showing the OS probability for patient that are stratified into macrophage engaged or excluded from based on their CD4^+^ T cell state

(G) Spearman correlations between patient survival and percentage of CD163 expressing CD163^+^/CD206^+^ macrophages, based on flow cytometry data

## DECLARATIONS

### Additional author information

ALB and FJB contributed equally. ALB and FJB are joint first authors.

### Contributors

ALB conceived, designed, and performed the experiments, analyzed and interpreted the results for the spectral flow cytometry data. FJB conceived, designed, and performed the experiments, analyzed and interpreted the results for the IMC data. DK, YV, and JW designed and supervised the research. ALB and FJB drafted the original manuscript, NB, JHWL, MMN, DK, YV, and JW reviewed and edited the manuscript. FH, MELV, MA, and SD performed the experiments and provided technical support. AB and NHV supported in data analysis. RRK assisted in patient inclusion and ROI selection. All authors read and approved the final manuscript. ChatGPT and Claude was used to improve readability and correct grammar.

### Ethics statements

Patient consent for publication

Written informed consent for this study was obtained for all patients, from themselves, parents or guardians.

### Ethics approval

Healthy adult donor blood was obtained from the shared healthy donor blood biobank from the University Medical Center Utrecht (biobank number 18-774). The institutional review board of the Erasmus Medical Center Rotterdam (MEC-2016-739) and the biobank committee of the Princess Máxima Center (PMCLAB2020.093) approved this study.

## Acknowledgements

We would like to acknowledge Elisa Zappa and Josephine G.M. Strijker for their help with the neuroblastoma sample preparation and Domenico Castigliego and Livio Kleij for their technical support.

