## Supplementary figures and images for "Inhibitory Nectin-2/TIGIT interactions between CD163^+^ macrophages and CD8^+^ T cells in spatially defined immune niches associate with poor outcomes in neuroblastoma"

### Supplementary figure 1

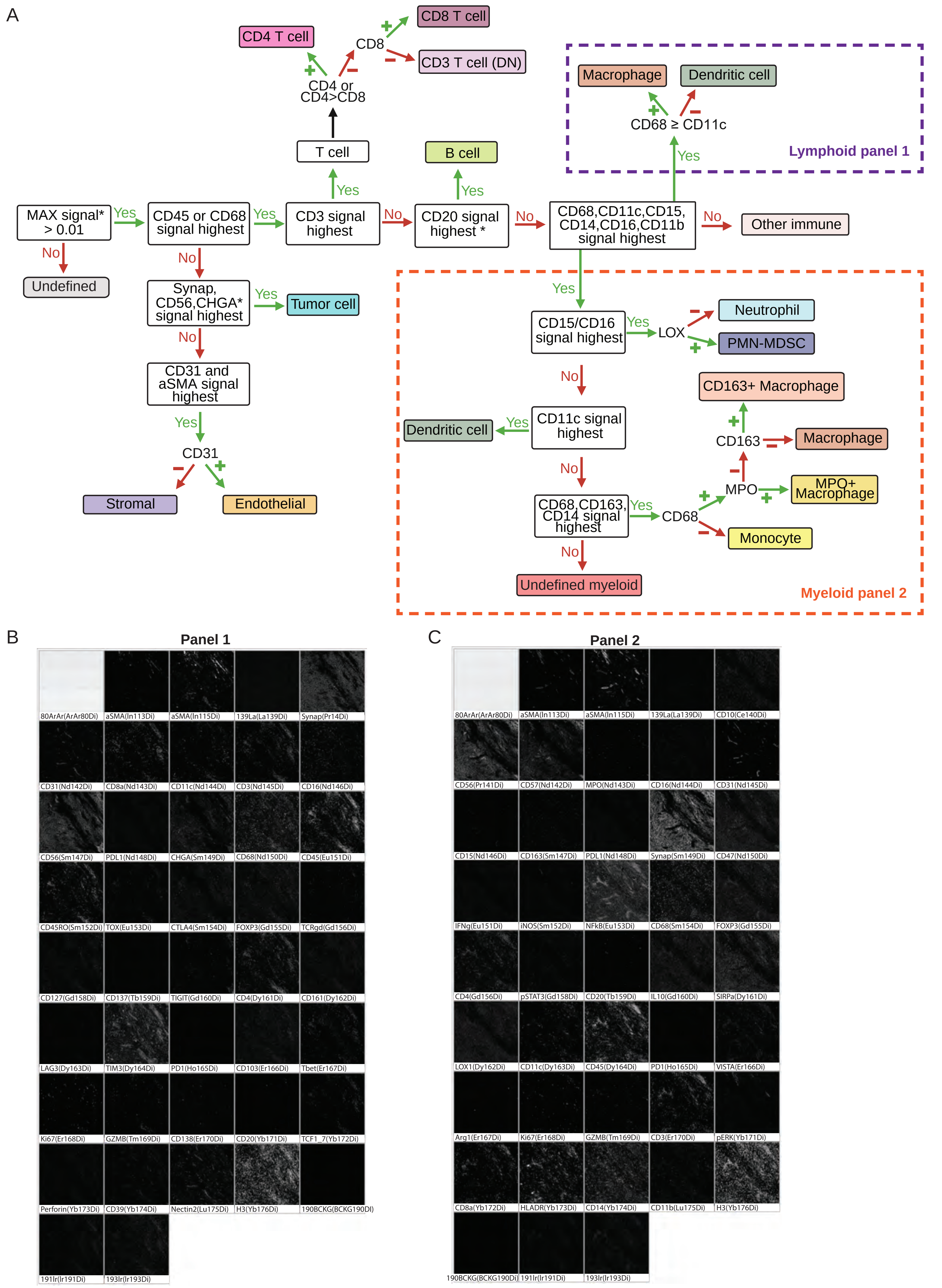

### Supplementary figure 2

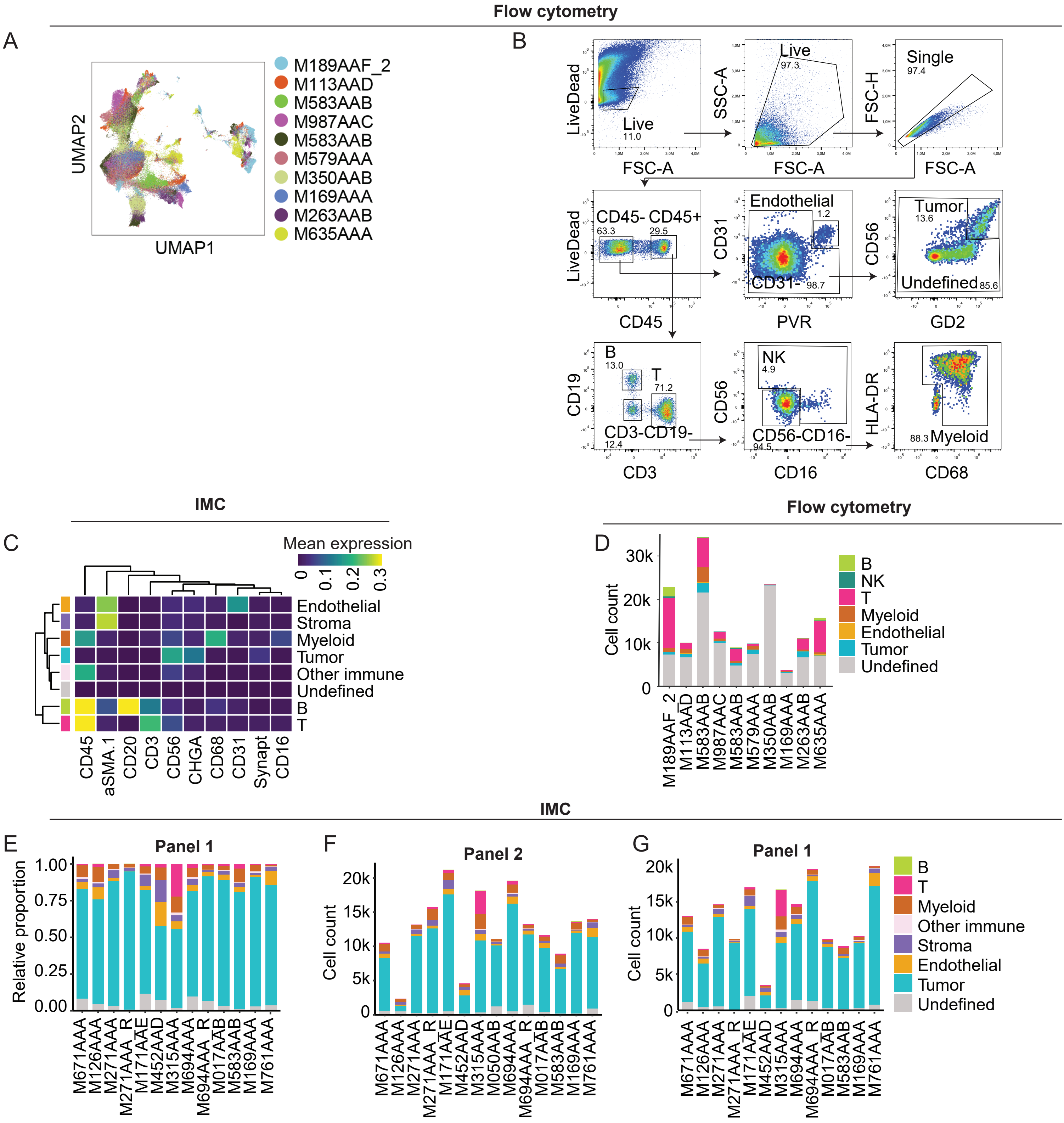

### Supplementary figure 3

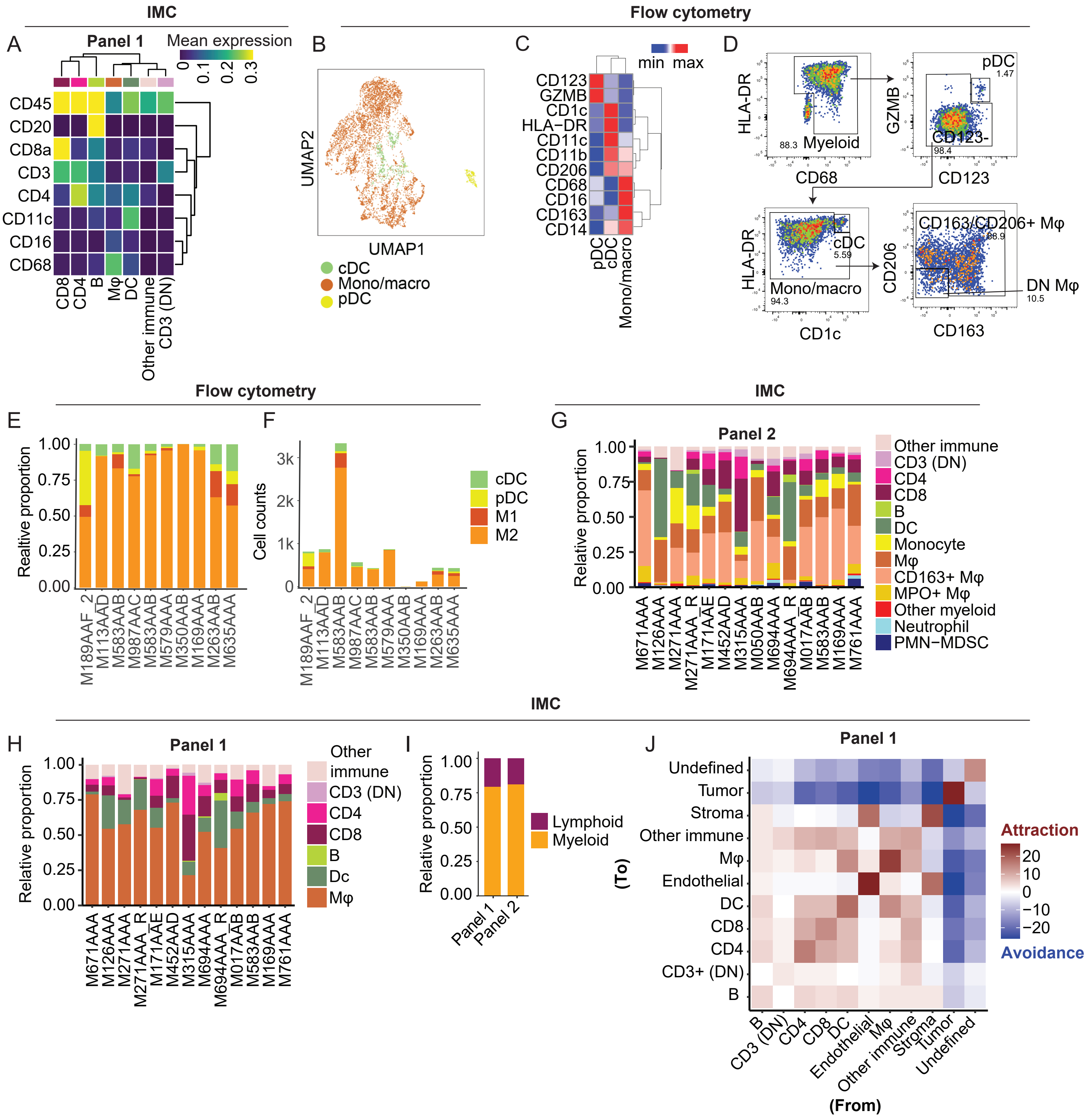

### Supplementary figure 4

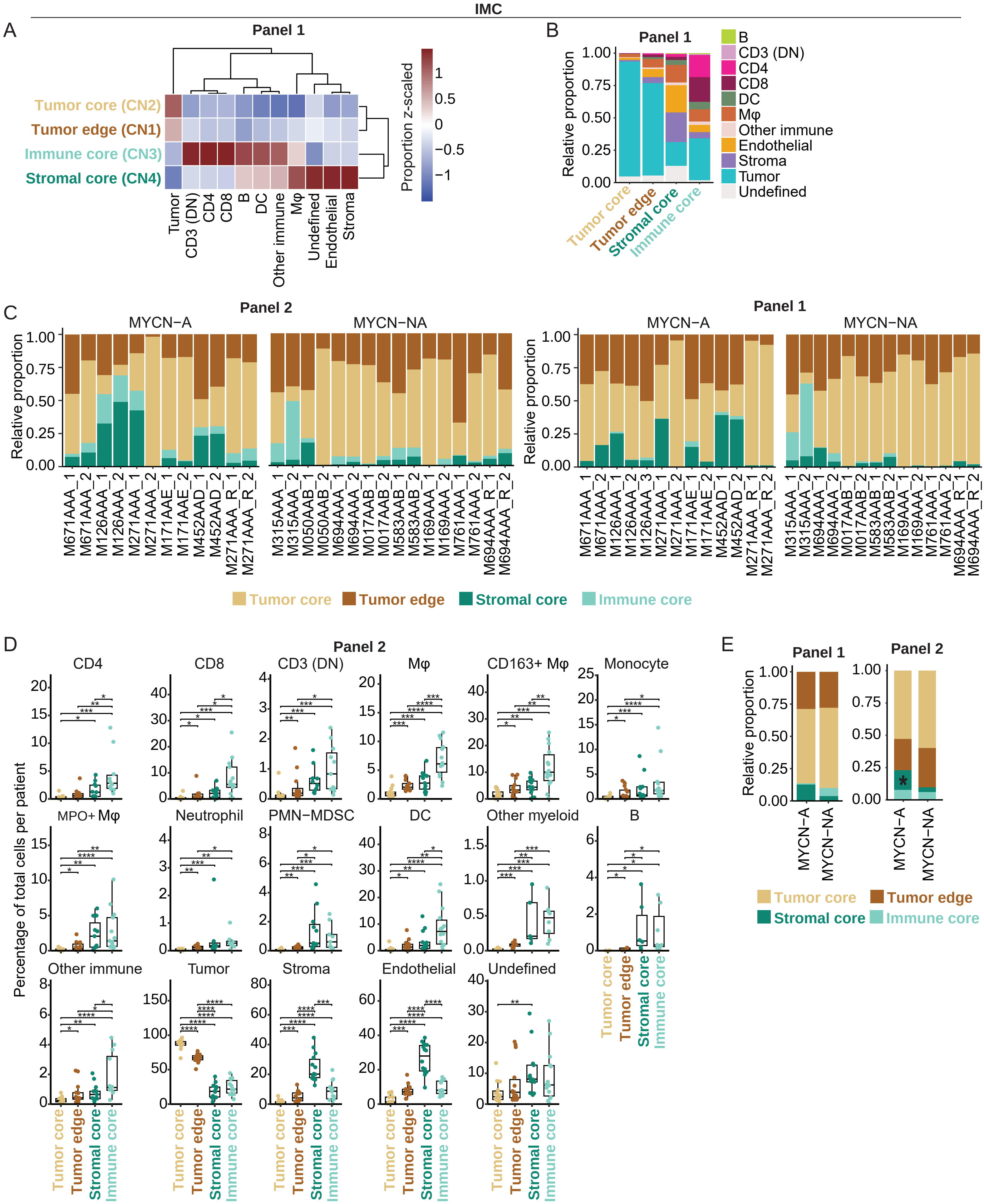

### Supplementary figure 5

Flow cytometry

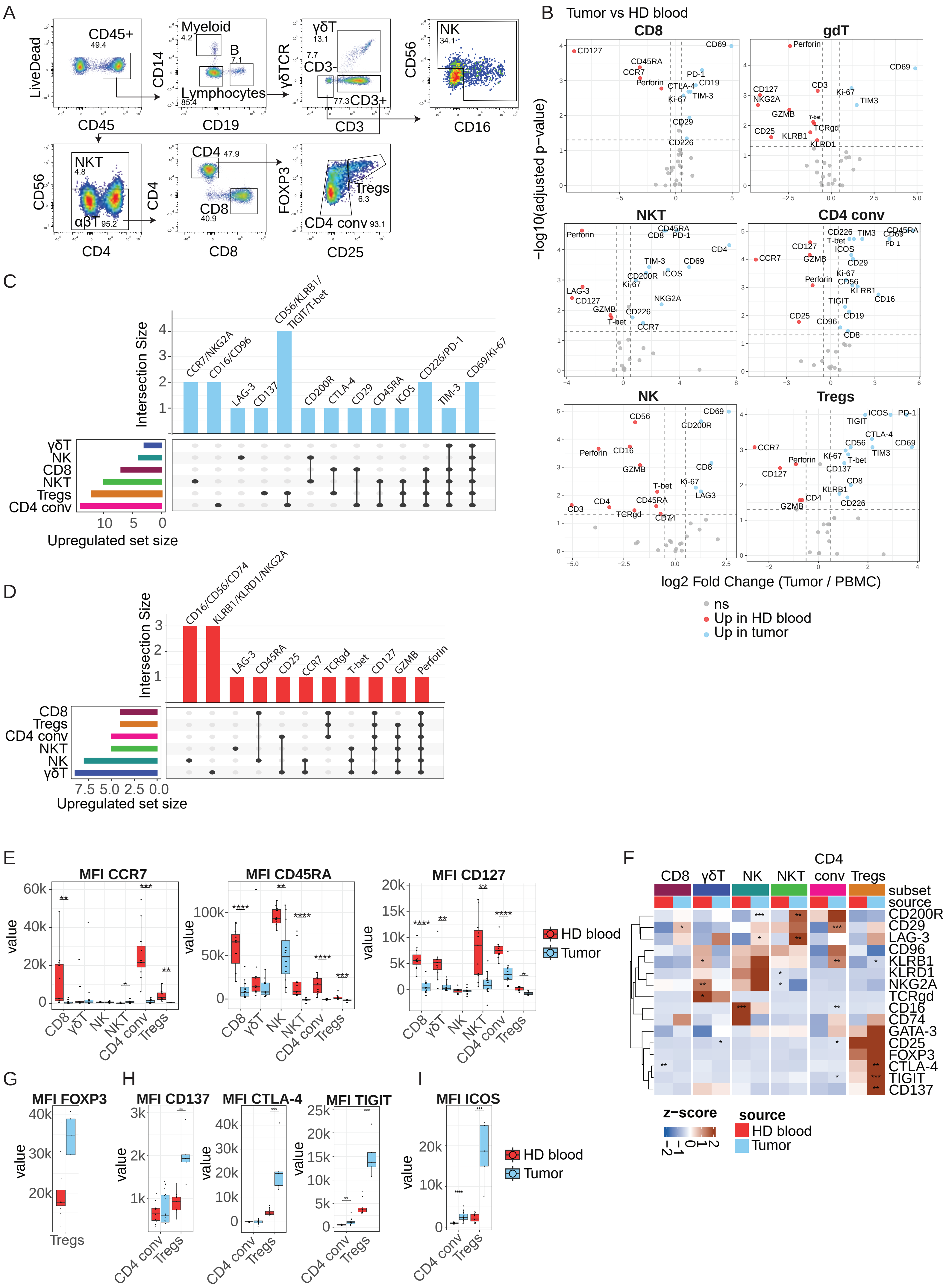

### Supplementary figure 6

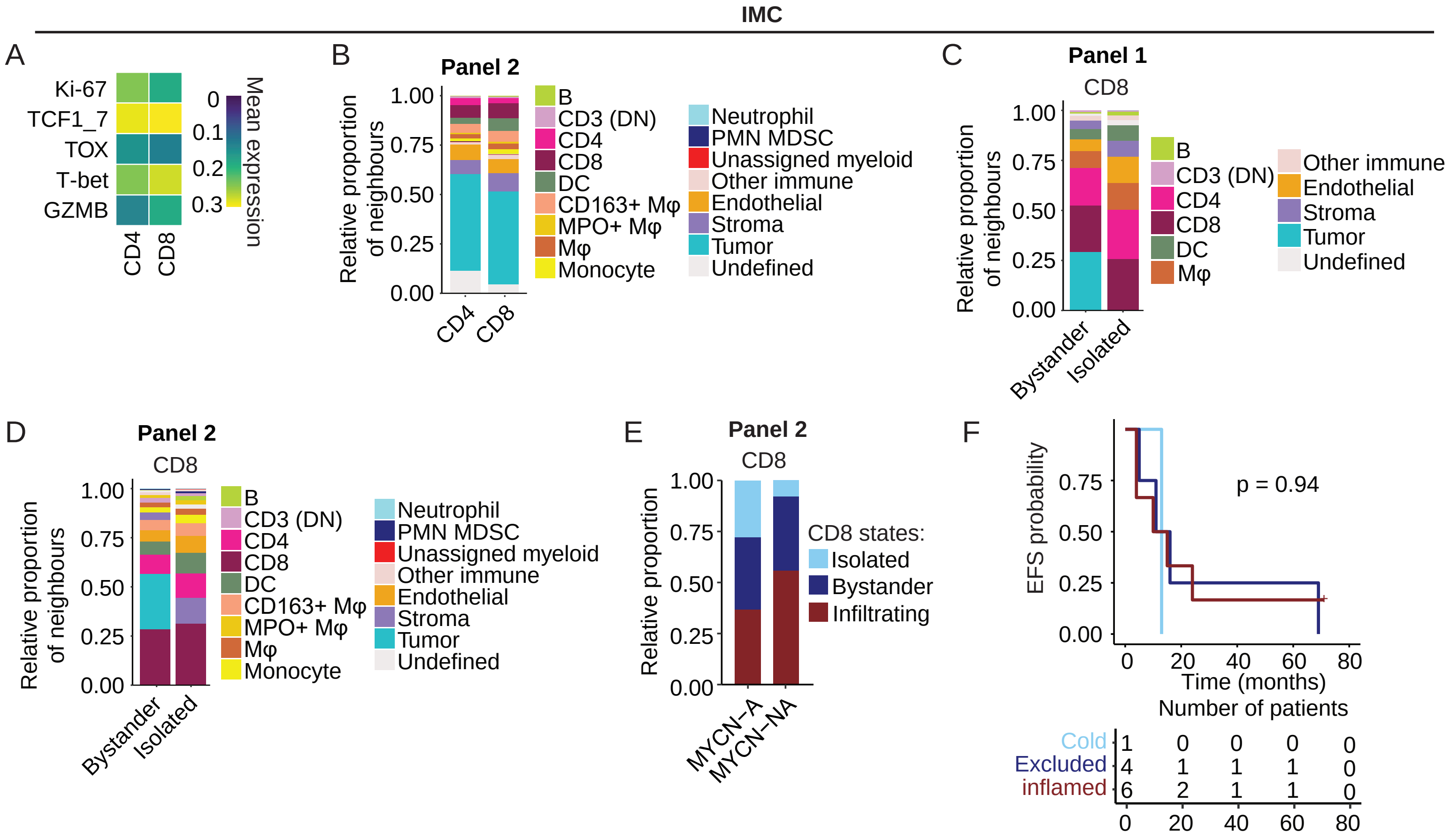

### Supplementary figure 7

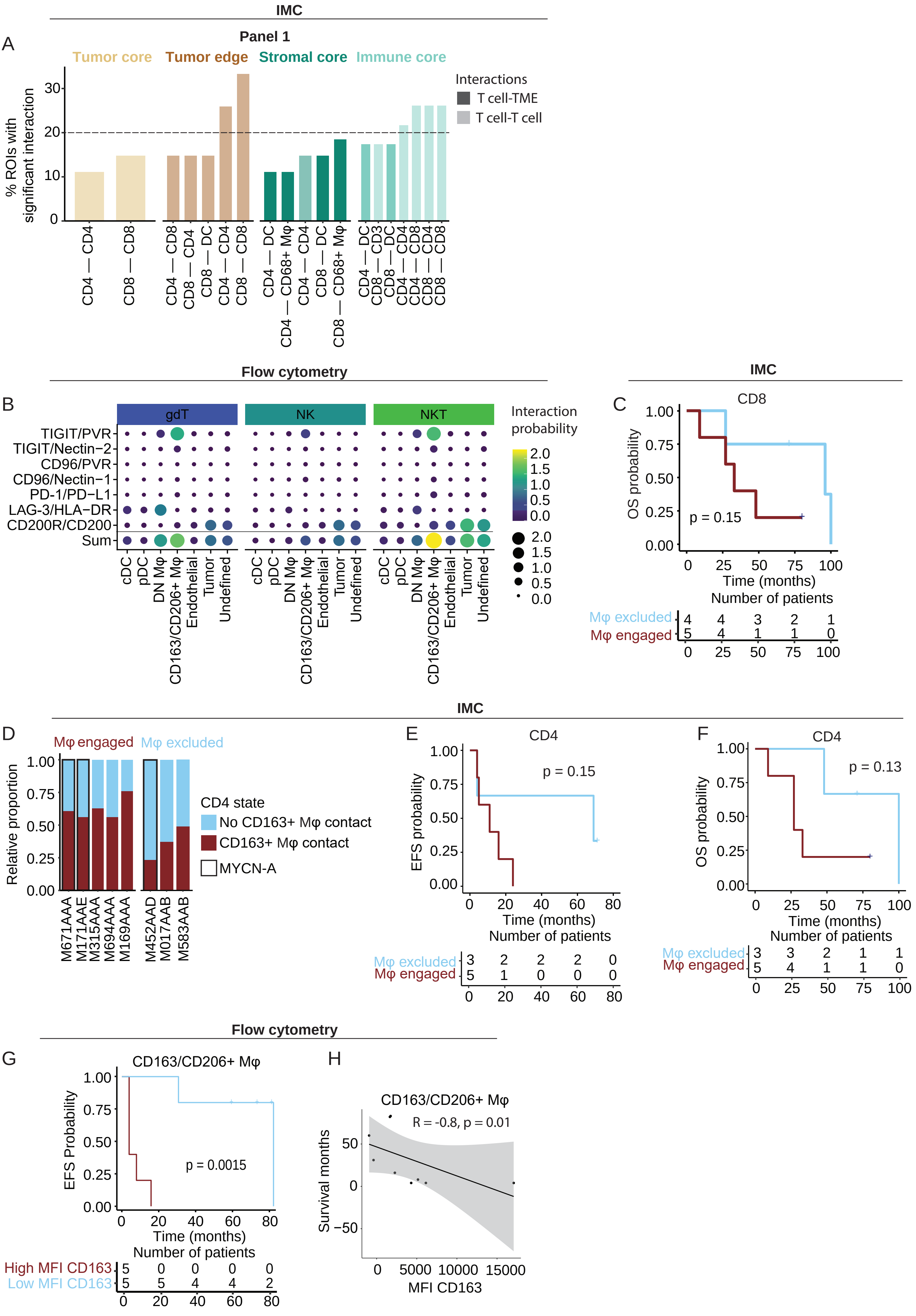
